# SNR Enhancement Considerations for Loop Receive Coils at Ultra-High Fields

**DOI:** 10.64898/2026.06.18.732714

**Authors:** Russell L. Lagore, Matt Waks, Timothy Hasapopoulos, Thomas Mercer, Andrea Grant, Yigitcan Eryaman, Kamil Ugurbil, Gregor Adriany, Alireza Sadeghi-Tarakameh

## Abstract

**Purpose:** To quantify parasitic losses in ultra-high field (UHF) magnetic resonance imaging (MRI) receive coils and determine how they contribute to the mismatch between numerically predicted and experimentally realized signal-to-noise ratio (SNR), with the goal of guiding receive-array designs toward ultimate intrinsic SNR (uiSNR).

**Methods:** SNR was measured across multiple field strengths (3T, 7T, 10.5T) using commercial and custom-built arrays. To quantify parasitic losses, unloaded-to-loaded quality factor ratio (Q_R_) measurements were performed on representative loop resonators and practical RF coils. Measured losses were combined with single-loop electromagnetic simulations to separate conductor, radiation, and component losses. These bench-derived loss estimates were then incorporated into full-array electromagnetic simulations of a 128-channel receive array to evaluate their impact on predicted intrinsic SNR.

**Results:** Measurements across field strengths supported the expected supralinear increase of SNR with B_0_. Q_R_ analysis showed that, at UHF, radiation loss must be excluded from unloaded-Q measurements to avoid overestimating electronic-noise penalties, and that multiple seemingly modest parasitic losses collectively impose substantial SNR degradation. In the 128-channel array, simulations including only conductor and radiation losses predicted 93% of central uiSNR, whereas inclusion of the full measured parasitic-loss budget reduced predicted performance to 78%, in close agreement with the experimentally measured 77%.

**Conclusions:** The gap between predicted and realized SNR performance of high-channel-count 10.5T loop arrays can be largely explained by parasitic losses that are not captured in conventional simulations. A bench-measurement-informed simulation framework enables more realistic prediction of coil performance and provides practical guidance for optimizing future UHF receive arrays.

## 1 Introduction

The demand for increasingly high spatial resolution in neuroimaging continues to grow, driven by the need to resolve fine-scale anatomical^1-4^ and functional features^5-7^ and to improve specificity in mapping brain circuits.^8-10^ However, higher resolution has typically been achieved at a fundamental cost, as scan time is increased and/or signal-to-noise ratio (SNR) is reduced. In practice, these constraints are tightly coupled: when higher SNR is available, it can be traded for higher resolution, shorter acquisition times, or both, enabling more efficient and robust imaging protocols. Ultra-high field (UHF; 7T and above) MRI is associated with inherently higher SNR compared with lower field strengths,^11-15^ providing a compelling pathway toward these goals. In this context, human imaging at 10.5T offers substantial potential for pushing spatial and/or temporal resolution beyond what is routinely feasible at 7T and below.^16-19^

Achievable SNR from increasing field strength is fundamentally bounded by the ultimate intrinsic SNR (uiSNR), which reflects physical limits imposed by the sample and electromagnetic field behavior.^20,21^ For neuroimaging applications, uiSNR as a function of field strength exhibits a characteristic spatial dependence, with near-quadratic trends in central regions^11,13,15^ and supra-linear behavior toward the periphery.^12,21^ Realizing these potential SNR gains depends critically on the performance of the radiofrequency (RF) receive (Rx) coil, which largely determines attainable SNR and the practical ability to exploit the intrinsic SNR benefits of UHF. Most modern head Rx coils are loop-based arrays, and increasing the loop count has traditionally been a reliable approach for improving SNR performance.^22-31^ However, maximizing SNR and parallel imaging performance requires careful optimization of array architecture, including the number of channels, element size, geometry, and, in some cases, element type. Within this framework, a practical and field strength agnostic way to assess Rx coil performance is to quantify how closely a given coil approaches the uiSNR limit, for example by using an SNR-to-uiSNR ratio.^32^

Electromagnetic simulation has been central to optimizing Rx arrays and predicting SNR performance within the uiSNR framework. Early numerical studies using idealized and/or generic coil models suggested that a large fraction (>80%) of uiSNR in intermediate-to-central regions could be captured with moderate-to-high channel-count (∼32-channel) loop arrays from 1.5T up to 9.4T.^33^ Subsequent work by Zhang et al.,^34^ incorporating coil-specific modeling and more realistic representations of physical arrays, demonstrated that at 7T a substantial fraction of uiSNR in intermediate-to-central regions is captured by 32- and 64-channel arrays, with coil-specific simulations predicting capture of 80%–90% and experimental measurements reporting 70%–80% of intermediate and central uiSNR. The observed ∼10% discrepancies between numerical and experimental results were initially attributed to unmodeled or difficult-to-quantify parasitic losses in practical systems.

At 10.5T, however, the same coil-specific modeling framework indicated a different trend. Even with 64 channels, the predicted fraction of intermediate-to-central uiSNR that could be captured was lower (on the order of 60%–70%), while experimental performance with a 64-channel Rx-only coil captured an even smaller fraction (∼40%–50%).^34^ More recent strategies have attempted to narrow this gap between the realized SNR and uiSNR. For instance, Waks et al.^31^ introduced an approach at 10.5T that leverages structures from a 16-channel transmit (Tx) array by incorporating them into a 64-channel Rx-only array to improve SNR, boosting the array’s central SNR performance from ∼50% to ∼70%. Lagore et al. combined this strategy with increased channel count (i.e., incorporating a 16-channel transceiver array into a 112-channel Rx-only array) and assembled a 128-channel Rx array, reporting capture of approximately ∼80% of central uiSNR, up from ∼70% with the Rx-only array.^30^ Despite these advances, the underlying reasons for persistent simulation–experiment mismatch^34^ at 10.5T—and the apparent difficulties of these initial high-channel-count traditional Rx-only arrays (64- and 112-channel) to consistently approach uiSNR^30,31^—remain poorly understood and motivated us to more systematically explore all the components in the receiver coil path. Here, a leading hypothesis is that unaccounted for parasitic losses - particular radiative losses become increasingly dominant at the ultra-high frequencies, yet prior work has not established a systematic, quantitative pathway to measure these losses, minimize them, and incorporate them into modeling in a way that explains the observed discrepancies.

A natural candidate for probing such losses is quality factor (Q) measurement—in particular, the unloaded-to-loaded Q ratio (Q_R_)—which is widely used to characterize loop coil losses.^35,36^ When accurately quantified, this metric provides a useful measure of how closely a practical coil approximates the performance of an ideal, lossless loop. However, conventional Q-measurement procedures do not necessarily capture the full set of parasitic losses present in practical multi-channel arrays, including losses associated with matching networks, detuning components, feed cables, and other implementation details that are challenging to model directly. Accordingly, measurement methodologies are needed that can quantify these loss channels in a manner that is both experimentally tractable and directly translatable to improved simulation accuracy.

In this study, we first perform a systematic experimental assessment of SNR across multiple field strengths (3T through 10.5T) and Rx coil implementations to establish the empirical trend of SNR gain with field strength and coil performance. We then use Q-factor measurement to quantify previously unaccounted parasitic losses relevant to practical coil implementations. Finally, we use the resulting loss estimates to correct simulation models and to explain, in a focused case study, the origin of simulation–experiment mismatch and the underperformance of a high-channel-count array (128 channels) at 10.5T. Together, these results provide a measurement-driven framework for reconciling simulated and experimental SNR at UHF and for guiding next-generation Rx coil design toward the uiSNR limit.

## 2 Methods

### 2.1 Experimental Assessment of Receive Array Performance

The purpose of this section is to systematically assess the SNR performance of several commercial and custom-built receive array coils across different field strengths—from 3T to 10.5T—in order to establish a general and translatable trend for SNR as a function of both field strength and coil design. In addition to characterizing absolute SNR behavior, the experimentally realized SNR is quantified relative to the ultimate intrinsic SNR limit, uiSNR. This normalization enables direct comparison across scanners and hardware using an SNR/uiSNR metric and helps identify the practical limitations and challenges that arise as coil arrays are pushed into the new ultra-high field regime.

#### 2.1.1 MRI Scanners

All MRI experiments were performed on whole-body scanners from Siemens Healthineers (Erlangen, Germany). Data were acquired on a 3T Prisma, 7T MAGNETOM, and 10.5T MAGNETOM. Across field strengths, the same general SNR measurement protocol, as well as post-processing technique, was used.

#### 2.1.2 3T Receiver Coils

At 3T, SNR measurements were performed using three commercially available receive arrays representative of commonly used head imaging configurations: a *20-channel head/neck coil, 32-channel head coil*, and a *64-channel head/neck coil* (all from Siemens Healthineers).

These coils span a range of channel counts and geometries and therefore provide a practical baseline for how SNR and SNR/uiSNR behave for standard clinical hardware at 3T.

#### 2.1.3 7T Receiver Coils

At 7T, SNR measurements were performed using two commercial arrays: a *32-channel head coil* (Nova Medical, Wilmington, MA) and a *64-channel head coil* (MR CoilTech, Glasgow, United Kingdom).

The 32-channel coil represents a widely used commercial configuration for high-resolution neuroimaging at 7T, whereas the 64-channel array reflects a state-of-the-art high channel-count design that has been engineered for enhanced SNR performance. Together, these coils allow evaluation of how increasing channel count at 7T impacts SNR and uiSNR capture under controlled conditions.

#### 2.1.4 10.5T Receiver Coils

At 10.5T, SNR measurements were performed using five custom-built receive array designs shown in Figure 1A-C: a *32-channel head coil*,^27^ a 64*-channel head coil*,^31^ an *80-channel head coil*,^31^ a *112-channel head coil*,^30^ and a *128-channel head coil*.^30^

**Figure 1:**
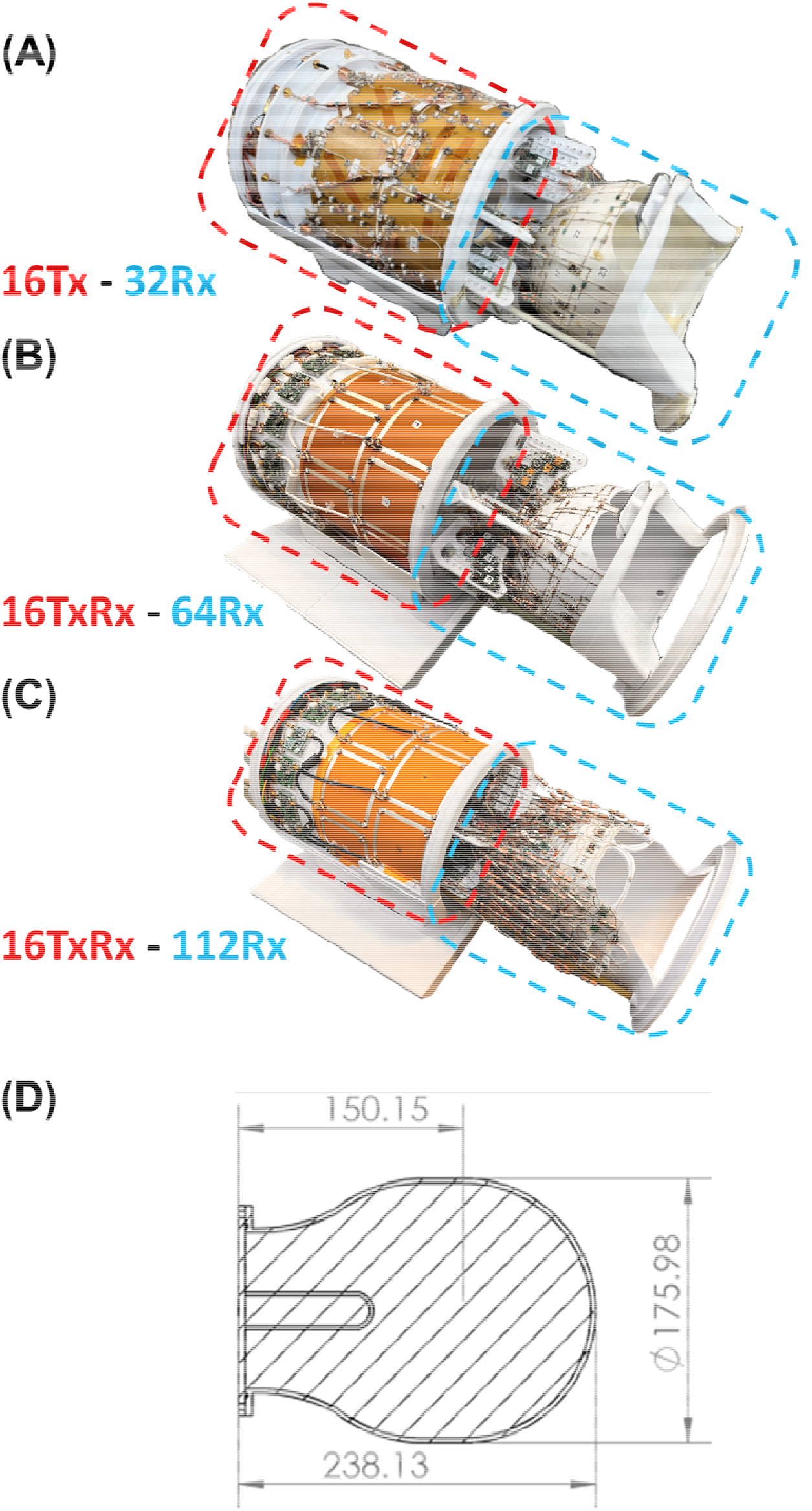
10.5T RF head coils studied in this work include a 16 transmit-only 32 receive only array (A), 16 transceive and 64 receive-only array (B), and 16 transceive and 112 receive-only array. A drawing of a light-bulb shape test phantom is shown in (D).

The 32-, 64-, and 112-channel arrays extend conventional loop-based architectures into the 10.5T regime, whereas the 80- and 128-channel arrays incorporate 16 large transceiver loop elements into the receive architecture. These hybrid Tx/Rx designs have been shown to improve central SNR,^30,31^ and are therefore included here to assess how such strategies influence realized SNR and SNR/uiSNR at 10.5T.

#### 2.1.5 SNR Measurements and Analysis

SNR measurements were performed inside a lightbulb-shaped phantom (Figure 1D) using fully-sampled 2D gradient-echo (GRE) acquisitions with parameters chosen to approximate fully relaxed conditions across field strengths. At all three field strengths, magnitude images for SNR estimation were acquired with a long repetition time (TR = 10 s), echo time TE = 3.48 ms, receiver bandwidth 87 kHz, and voxel size 2.0 × 1.0 × 2.0 mm^3^.

Noise images were acquired using the same readout and receiver settings but with RF excitation disabled (nominal flip angle = 0°) and a reduced TR of 600 ms to shorten acquisition time. SNR maps were then constructed from the complex signal and noise data using the covariance-weighted root-sum-of-squares (cov-rSoS) approach,^37-39^ as previously described.^26,31^

All measurements were performed using a polyvinylpyrrolidone (PVP)–based uniform light bulb– shaped phantom,^27^ which approximates the size and loading of the human head and provides a stable, reproducible medium for cross-coil and cross-field-strength comparisons. The electrical properties of the phantom were measured at each field strength using a commercial dielectric assessment kit (DAKS-12, SPEAG, Zurich, Switzerland) and characterized as follows: at 3T, ε = 55 and σ = 0.47 S/m; at 7T, ε = 51 and σ = 0.56 S/m; and at 10.5T, ε = 48 and σ = 0.65 S/m.

The uiSNR was numerically calculated as previously described^34^ and used as a reference metric to assess coil performance across field strengths. To enable comparison with uiSNR, the intrinsic SNR was obtained by scaling the experimental SNR values to remove the effects of sequence parameters, flip angle, and phantom properties, following the formalism described in literature^32,29,31^ and summarized in Eq. (1):

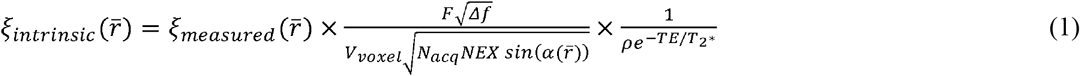

where 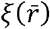 is the SNR at location 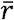, *F* is the noise factor of the system (assumed to be 1.15 across all systems), Δ*f* is the receiver bandwidth, *V*_*voxel*_ is the voxel volume, *N*_*acq*_ is number of the acquired *k*-space samples, *NEX* is the number of averaging, 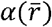 is the measured flip angle at location 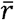, *ρ* is the proton density of the PVP phantom (measured as 0.69), and *TE* is the echo time. The 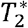 of the phantom was measured as 24ms at 10.5T and linearly extrapolated to 36 ms and 84 ms at 7T and 3T, respectively.

To evaluate SNR performance as a function of depth from the surface of the phantom, three-dimensional SNR maps were analyzed using light bulb–shaped shell regions of interest (ROIs) concentric with the phantom. Shell ROIs with a thickness of 1 cm were defined at multiple depths from the outer phantom surface toward the center, and SNR was averaged within each shell to obtain depth-dependent SNR profiles. Central SNR was quantified by averaging within a spherical ROI with a radius of 1 cm located at the geometric center of the phantom.

### 2.2 Quantification of Parasitic Losses

The ohmic losses that contribute to the thermal noise of a receive coil can be broadly divided into two categories: sample loss and parasitic loss.^35^ Parasitic losses degrade the SNR of an Rx coil by introducing additional thermal noise sources and, for a generic Rx coil, can be attributed to several components, including coil loss (a combination of conductor and substrate losses), radiation loss, lossy lumped elements used primarily for coil tuning (mainly capacitors), lossy matching networks (typically comprising capacitors and inductors), and lossy detuning networks (typically comprising a PIN diode and a tank circuit). In this study, these individual loss components were estimated using a hybrid experimental–computational Q_R_–based technique.

Experimental quantification of parasitic losses was achieved through Q-factor measurements of various RF loop coils on the bench. Losses were broken down into various categories through iterative measurements of resonant structures with progressively more circuitry added, going from the bare resonator all the way through to addition of feed circuitry and cabling.

#### 2.2.1 Probe Construction

A double decoupled B-field probe^40^ was used for all bench measurements of Q-factor. It is constructed by overlapping a pair of shielded loop B-field probes to achieve geometric decoupling between the probes. An adjustable probe was constructed as depicted in Figure 2A. The probe is constructed from UT-141C (Amphenol CIT, St. Augustine, FL, USA) coaxial cable connectorized with bulkhead BNC connectors for connection to the vector network analyzer (VNA). It features an adjustment screw to vary the overlap between the probes. While the probes are reasonably well decoupled over a wide bandwidth, this adjustment screw can be used to optimize decoupling to < -60 dB at a frequency of interest.

**Figure 2:**
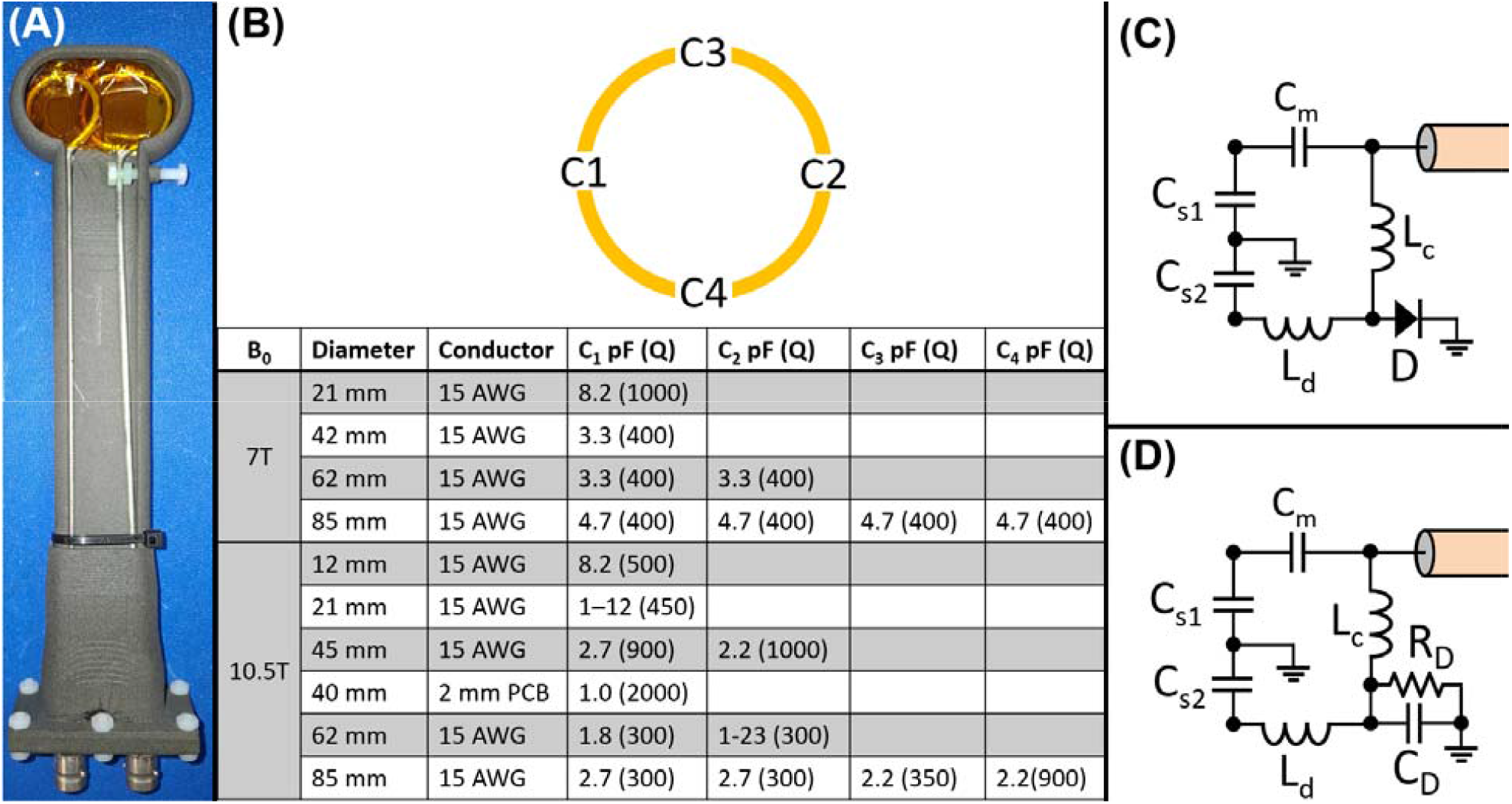
(A) A double-decoupled magnetic field probe used for Q-ratio measurements. This probe features an adjustment screw which allows for optimal decoupling between the probes. contains details on the resonators which were studied with respect to Q-ratio as sample distance increases. For practical RF coils measured, the feed circuit shown in (C) as used with its equivalent circuit shown in (D).

#### 2.2.2 Resonator Construction

To verify the validity of the simulations, a set of Q_R_ measurements was first performed on simple loop resonators without matching or detuning networks, spanning a range of loop sizes and coil-sample distances (1 cm to 5 cm). The investigated resonators were circular loops with diameters of 2 cm, 4 cm, 6 cm, and 8 cm, implemented either as printed circuit board loops or as copper-wire loops. These resonators were tuned for 7T (297 MHz) and 10.5T (447 MHz). Details are listed in Figure 2B. Resonators contain one to four segmenting capacitors (C1 - C4) with capacitance values and Q-factors also listed in Figure 2B. Q-factors for these components, and thus the relevant ESRs, were derived from the manufacturer datasheets.

#### 2.2.3 RF Coil Construction

A loop resonator intended to be used as an RF coil needs additional components. Practically, these loop resonators need components for impedance matching and detuning (in the case of receive-only coils). A sub-set of the resonators described in 2.2.2 (2 cm, 4 cm, and 6 cm diameter copper wire loops) were constructed with inclusion of one type of feed circuit depicted in Figure 2C. An equivalent circuit showing parasitic and dominant losses of this feed circuit is also shown in Figure 2D.

#### 2.2.4 Q-factor measurement

Q-factors are measured with the probe described in 2.2.1 using a transmission (S_21_) 3 dB bandwidth measurement method on a 16-port VNA (ZNBT8, Rohde & Schwarz, Germany). Q-factor is calculated simply as:

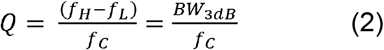

The VNA source power is set to +10 dBm and frequency span is adjusted such that the resonant peak fills a large part of the measurement. A minimum of 201 points are used in the sweep, but often 501 and 1001 points are used, particularly for very high Q resonators, to resolve the peak accurately. The measurement bandwidth is reduced appropriately to lower the instrument noise floor adequately while still being fast enough to make practical measurements.

The probe is held high above the resonator being measured and approached slowly until a clean spectrum is acquired above the noise floor. This measurement must be made at the furthest practical distance that produces a reproducible measurement, since the probe will introduce losses to the resonator and consequently reduce Q if it is brought excessively close. This distance depends upon the size of resonator. Larger resonators create larger areas of sensitivity than small resonators.

#### 2.2.5 Unloaded Q measurement

Unloaded Q (Q_u_) is a measurement of the ratio of coil inductive reactance (X_Coil_) to coil resistance (R_Coil_) . In the case of a bare resonator, R_Coil_ is composed of capacitor losses (R_Cap_), conductor losses (R_Cond_), and radiation losses (R_Rad_) whereas a coil with feed and detune circuitry also adds match and detune parasitic losses (R_Match_ and R_Detune_, respectively).

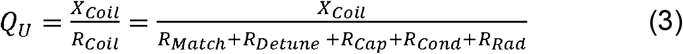

Because this work is focused on ultra-high field, resonant loops are treated like antennas which have a significant component of R_Rad_ which must be isolated and quantified separately. Consequently, unloaded Q measurements are made as described in Chen et al.^41^ in proximity to a conductive copper sheet. This method was proposed for antenna Q measurements where radiation resistance (R_Rad_) is high, but is necessary for UHF loop resonators due to the high R_Rad_ of unloaded loops. The distance from the conductive sheet is incremented with foam padding from 1 cm up to 5 cm to maximize Q_U_, thus minimizing or eliminating R_Rad_ from the measurement.

#### 2.2.6 Loaded Q measurement

Loaded Q (Q_L_) is a measure of sample resistance (R_Sample_) and R_Coil._ In the loaded case, R_Rad_ is presumed to be negligible and R_Coil_ is known, so R_Sample_ can be determined from measurement of Q_L_.

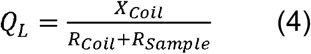

The measurement is made with the same techniques as before, but it is measured with the resonator or coil in proximity to a cube phantom 168 mm on each side containing a solution with ε_r_=78 and σ=0.95 - 0.97 S/m (300 - 450 MHz) as measured with a SPEAG DAKS-12 dielectric probe. Distance of the resonator to the load is incremented by 1 cm using foam padding, with the first measurement at 1 cm placed directly against the 1 cm acrylic wall of the phantom.

#### 2.2.7 Q-ratio and SNR calculations

Unloaded and loaded Q factors are measured for the principal purpose of determining the performance of the coil. A ratio of these quantities is the Q-ratio (Q_R_ = Q_U_/Q_L_) which is a useful metric describing the ratio of electronic to sample noise. The resultant SNR loss as a result of electronic sources of noise can be calculated^42^ as:

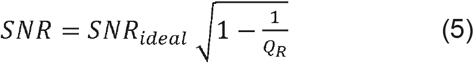

#### 2.2.8 Estimation of Individual Parasitic Losses

Simulations were performed using a frequency-domain finite-element solver (HFSS, ANSYS, Canonsburg, PA). The loops were modeled either as copper wires with their realistic thickness or as planar loops printed on an FR4 substrate, consistent with the corresponding physical elements. A lumped voltage port with a 50Ω characteristic impedance was assigned, and the reflection coefficient (*S*_11_) was calculated as a function of the applied input voltage. Distributed tuning capacitors were placed at representative locations on the loops using their nominal bench values and equivalent series resistance (ESR) values from the manufacturer datasheets. Each loop was excited with a 1 V forward voltage, which was translated into the total current at the port using the following expression:

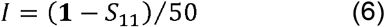

To estimate the resistances associated with the sample *(R*_*Sample)*_, conductor *(R*_*Cond*_), and radiation *(R*_*Rad*_), the corresponding power losses *(P)* were exported separately and normalized to the input current as follows:

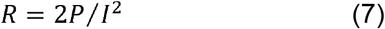

The Q_R_ of the simulated resonant loops was then calculated as:

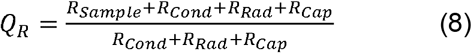

Once good agreement between simulation and experiment was achieved, the framework for the Q measurement and simulation was established, and Q-ratio trend was obtained at different ultrahigh field strengths (i.e., 7T and 10.5T)

Next, this whole process was repeated for four different loop geometries—4 cm–diameter circular, 5×5 cm^2^ square, 7×4 cm^2^ elliptical, and 4 cm equilateral triangular—each representative of different loops on the physical 128-channel array at 10.5T. These measurements were performed at four distances from the phantom surface (1–4 cm in 1 cm increments), representative of realistic loading conditions. After good agreement between simulation and experiment was achieved, and the simulated sample, conductor, radiation, and capacitor resistances were thereby validated, the losses associated with the remaining Rx loop circuitry—namely the matching and detuning networks—were estimated by incremental bench measurement of Q_R_. To do this, the matching network was first added to the resonant loop and updated QR was measured. The loss associated with the matching network *(R*_*Match*_) was then obtained by solving the following equation, with all other parameters fixed from the validated simulations:

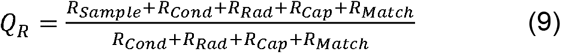

After *R*_*Match*_ was determined, the detuning network was added as the final component of the Rx loop circuitry. The updated Q_R_ was then measured, and the detuning-network loss *R*_*Detune*_ was obtained by solving:

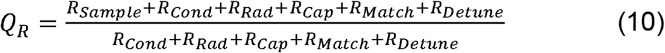

The sum of all non-sample losses was defined as the total parasitic loss:

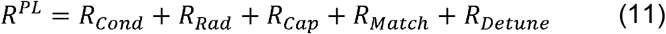

### 2.3 Bench-Measurement-Informed Receive Array Simulation

This section describes how the 128-channel receive array was modeled, simulated, and post-processed. It also details how parasitic losses of the individual loops—quantified via bench measurements of the Q-ratio as described in Section 2.2—were incorporated into the simulation framework to adjust the simulated SNR and more accurately reflect the performance of the physical array.

#### 2.3.1 EM simulation of the Receive Array

The major portion of the 128-channel coil was constructed by wrapping a thin flexible PCB (FPC) around the coil former. The exact geometry of these FPC-based loops was therefore available within SolidWorks (Dassault Systèmes, Vélizy-Villacoublay, FR). While vias and a bottom layer were employed to jumper loops, this geometry could not be realized in CAD, so jumper wires were added to represent the overlapping segments of the loops. Loops without a corresponding FPC layout (mainly those placed on the former by hand) were manually modeled to replicate the real-life conductor paths as closely as possible. The reconstructed conductors in SolidWorks were then placed on a polyethylene former, and the light-bulb phantom model was positioned inside the former to match the geometry and positioning used in the MRI experiments. The complete assembly was exported as a STEP file and imported into the electromagnetic (EM) simulation environment (HFSS, ANSYS, Canonsburg, PA), as illustrated in Figure 3.

**Figure 3:**
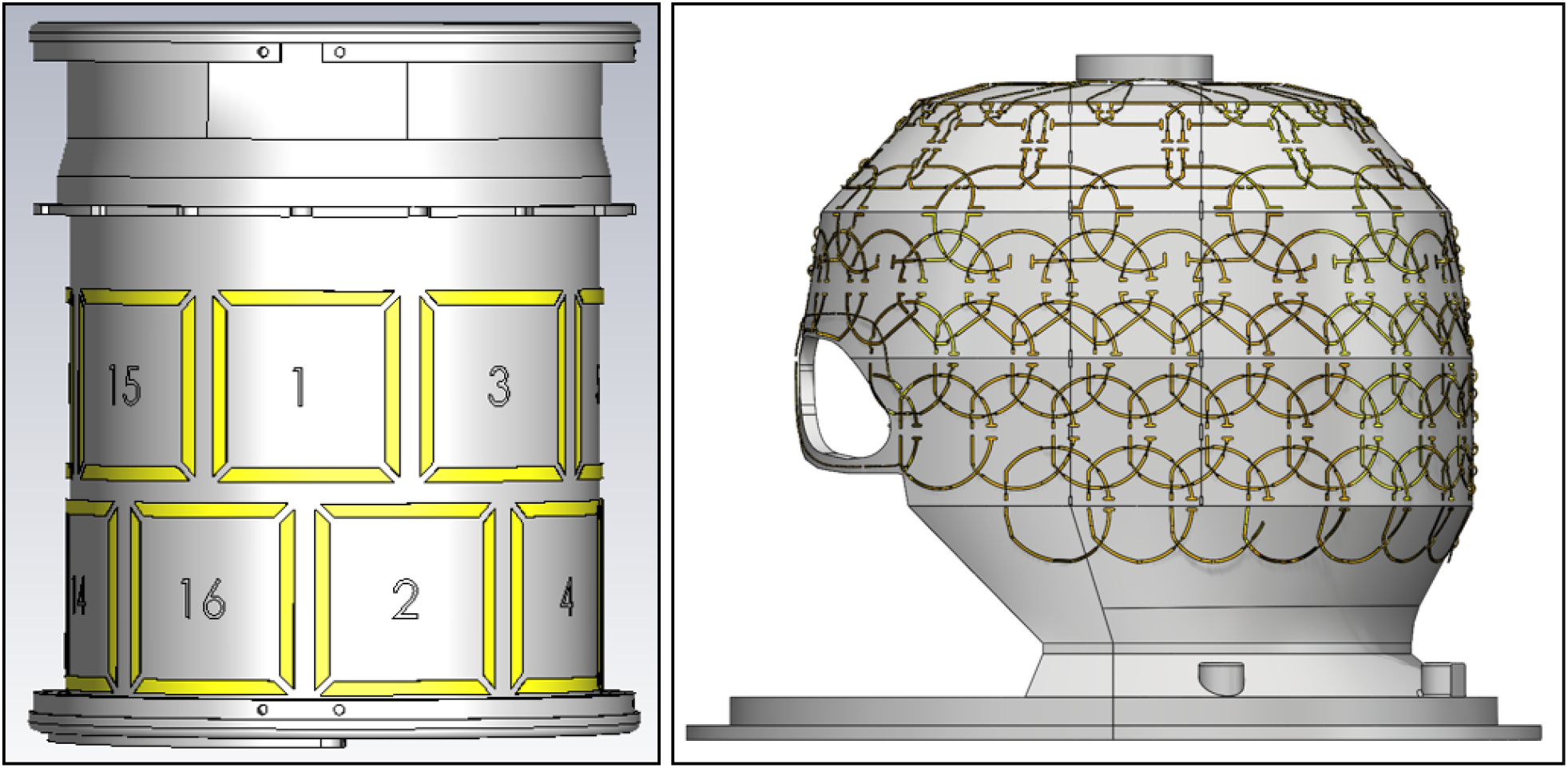
The complete transceiver (left) and receiver (right) CAD assemblies of the 128Rx used for electromagnetic (EM) simulations.

In the EM simulations, HFSS was configured in “*Network Analysis”* mode with the “*Terminal”* solution type. All conductors were modeled as silver, and all capacitors on the loops were assigned their nominal values from the physical setup. No explicit matching circuitry was modeled. Instead, all excitation ports were assigned as “*Terminal Lumped Ports”* with a 3 kΩ characteristic impedance to emulate the preamplifier decoupling used in the real configuration. A “*Radiation”* boundary condition was applied around the model using the HFSS “*Auto Region”* setup to ensure negligible reflection of EM waves from the outer boundaries. An “*Adaptive Mesh Refinement”* strategy was used with a single-frequency simulation setup. The “*Convergence”* criterion was set to 10^−3^ on the S-parameters, with a maximum of 20 adaptive iterations. All simulations were performed on a workstation equipped with two 8-core Intel® Xeon® W-2245 processors (3.9 GHz) and 256 GB RAM.

#### 2.3.2 Intrinsic SNR Reconstruction

Once the simulation reached the convergence criterion, the last adaptive solution was used to export two sets of quantities necessary for SNR calculations: the left-handed circularly-polarized magnetic field 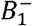 (representative of the receive sensitivity) and the S-matrix (S).

The 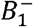 fields were exported on a per-channel basis by setting the *“Incident Voltage”* (*V*^+^) at the excitation port of the intended channel to 1 V, and 0 V for all other channels. Here, 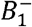 was defined as

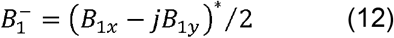

The full *N × N* S-matrix was exported into MATLAB and used with a characteristic impedance *Z*_*o*_ = 3 *k Ω*. to compute the full impedance matrix **Z** as

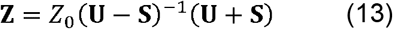

where U is the *N × N* identity matrix (i.e., *N* is the total number of channels). The same characteristic impedance was used to compute the incident current, (*I*^+^), at the excitation ports corresponding to the 1 V incident voltage, (*V*^+^), used for the 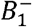 export as

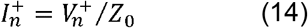

These incident currents were later used to normalize the sensitivity of each loop to unit current, as originally theorized by Roemer et al.^37^ The current-normalized sensitivity of the *n*^*th*^ loop at location 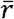 was calculated as:^32^

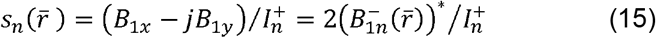

where 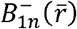 corresponds to the *n*^*th*^ loop at location 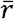.

The 1 *x N* complex vector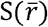

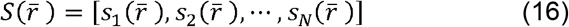

containing the sensitivities of all loops at location 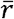 was then used with the cov-rSoS approach^39^ to compute the optimal intrinsic SNR^32^ (iSNR) as:

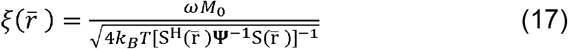

where *k*_*B*_ is Boltzmann’s constant, *T* is the absolute temperature of the sample (assumed to be 293 K), *ω* is the angular Larmor frequency at 10.5T, *M*_*o*_ is the equilibrium magnetization for hydrogen (calculated as 3.46×10^−2^ A.m^−1^ using the expression in^32^), and **Ψ** denotes the *N × N* noise covariance matrix. The matrix **Ψ** represents the thermal noise (ohmic loss) experienced by each loop, as well as the shared thermal noise between loops.

For current-normalized simulations, **Ψ** can be calculated from the **S** matrix using the expression introduced by Bosma et al.^43^ and recently revisited,^44^ reformulated here as follows:

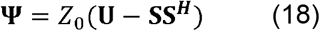

where ***H*** denotes the Hermitian operator (i.e., complex conjugate transpose). To account for all parasitic losses, the parasitic loss resistance measured for each loop (see Section 2.2) was added to the corresponding diagonal element of the **Z** matrix. In other words, elements of the corrected **Z** matrix were defined as

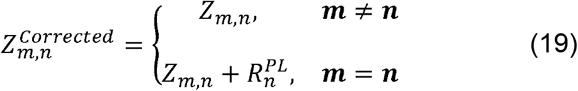

where 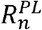 is the parasitic loss resistance of the *n*^*th*^ loop, as defined in Equation 11 and obtained following the procedure describe in Section 2.2.8 for representative loops of the 128-channel coil in different sizes, shapes, and loading distances. Loops in the simulation model with the closest match in size, shape, and loading distance to the measured loops were then assigned the corresponding *R*^*PL*^ values. Note that, for adjustment of the **Z** matrix, the *R*^*PL*^ values were used after excluding *R*_*Rad*_. The rationale for this exclusion was that radiation loss was already inherently accounted for in the simulation environment, unlike the other parasitic loss components. The corrected **Z** matrix was then used to updated the **S** matrix as follows:

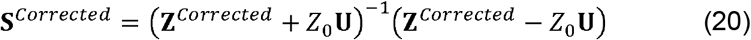

The corrected ***S*** matrix was then substituted into Equation 18 to calculate the corrected, which was subsequently used to calculate the intrinsic SNR in Equation 17.

The resulting intrinsic SNR inside the lightbulb-shaped phantom for the 128-channel coil was finally compared with the experimentally acquired iSNR, as well as with the uiSNR.

## 3 Results

### 3.1 Experimental Assessment of Receive Arrays Performance

Figure 4 illustrates axial slices of the intrinsic SNR maps for the 3T coils, with the left column showing the measured intrinsic SNR and the right column showing the same maps normalized by the uiSNR map. This normalization highlights how efficiently each coil configuration approaches the theoretical SNR limit at 3T.

**Figure 4:**
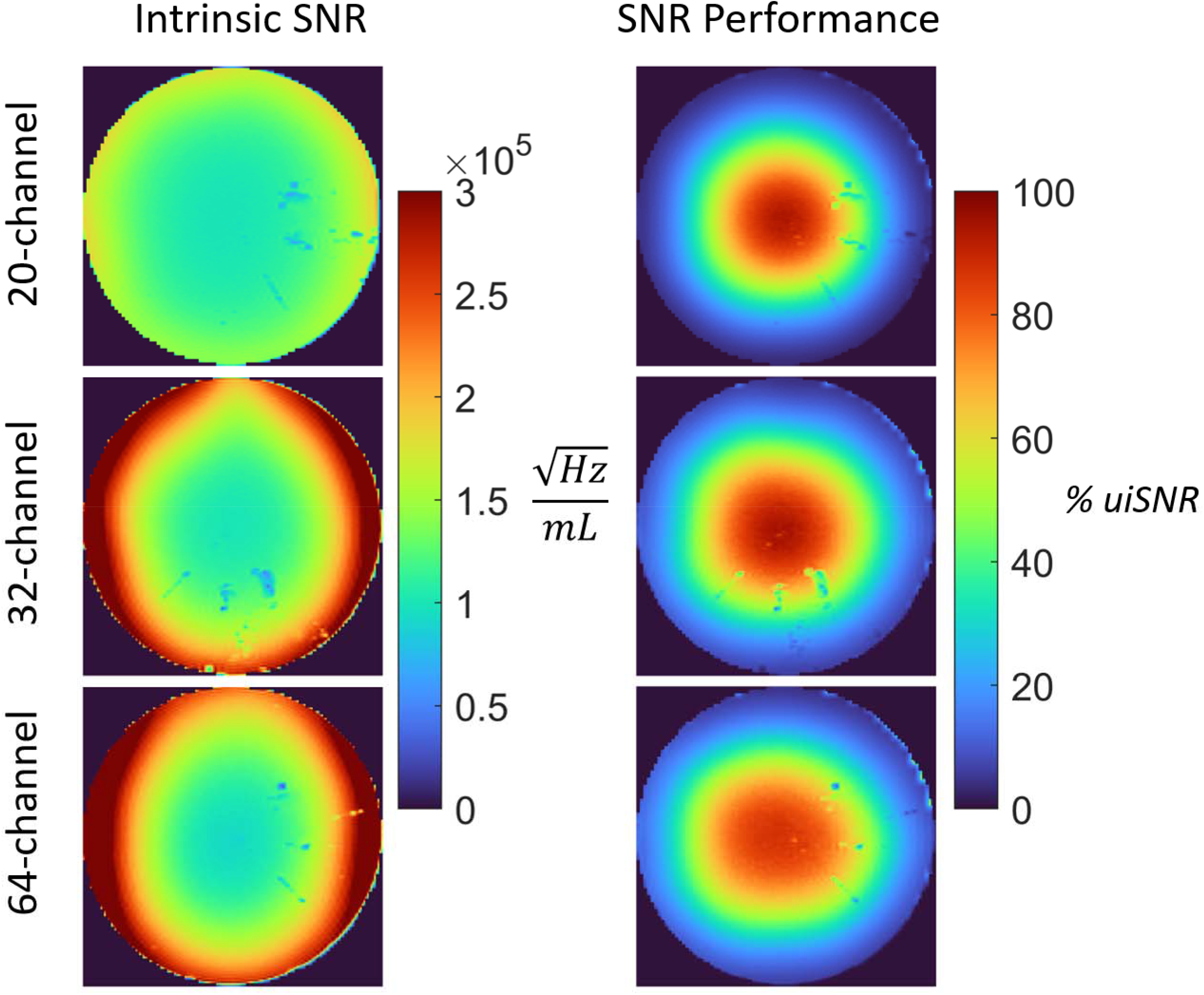
Axial slices of the intrinsic SNR maps (left column) for 20-channel, 32-channel, and 64-channel 3T coils. Also shown are iSNR maps normalized by the uiSNR map and expressed as a percentage of uiSNR.

Similarly, Figure 5 and Figure 6 present coil performance maps at 7T and 10.5T, respectively. For each field strength, the measured intrinsic SNR maps are displayed alongside their uiSNR-normalized counterparts, enabling direct visual comparison of how coil design and channel count influence both the magnitude and spatial distribution of SNR, as well as the fraction of uiSNR that is realized in central and peripheral regions.

**Figure 5:**
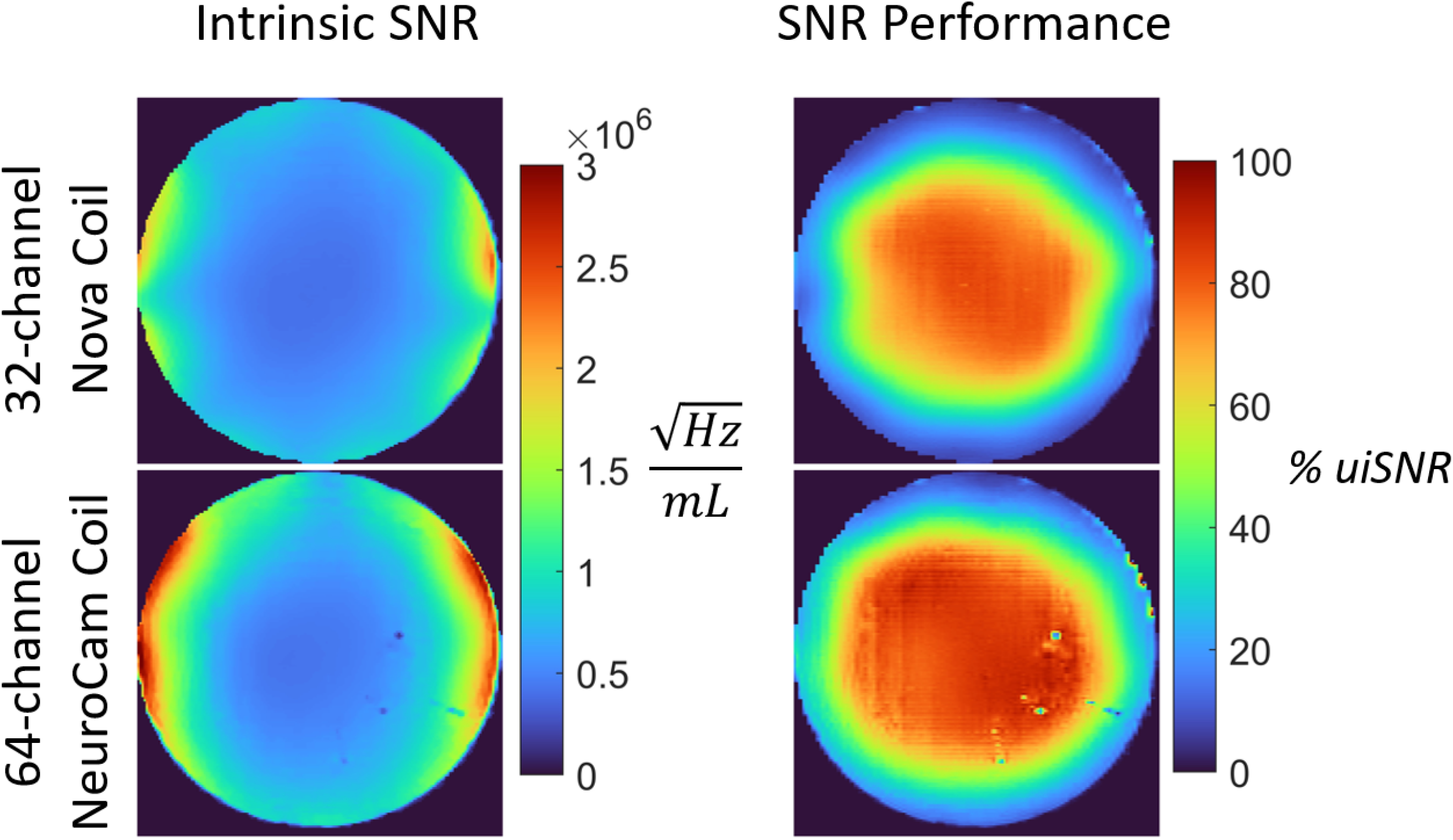
Intrinsic SNR maps (left) and uiSNR-normalized maps (right) for a 32-channel (Nova) and 64-channel (NeuroCam) coil at 7T.

**Figure 6:**
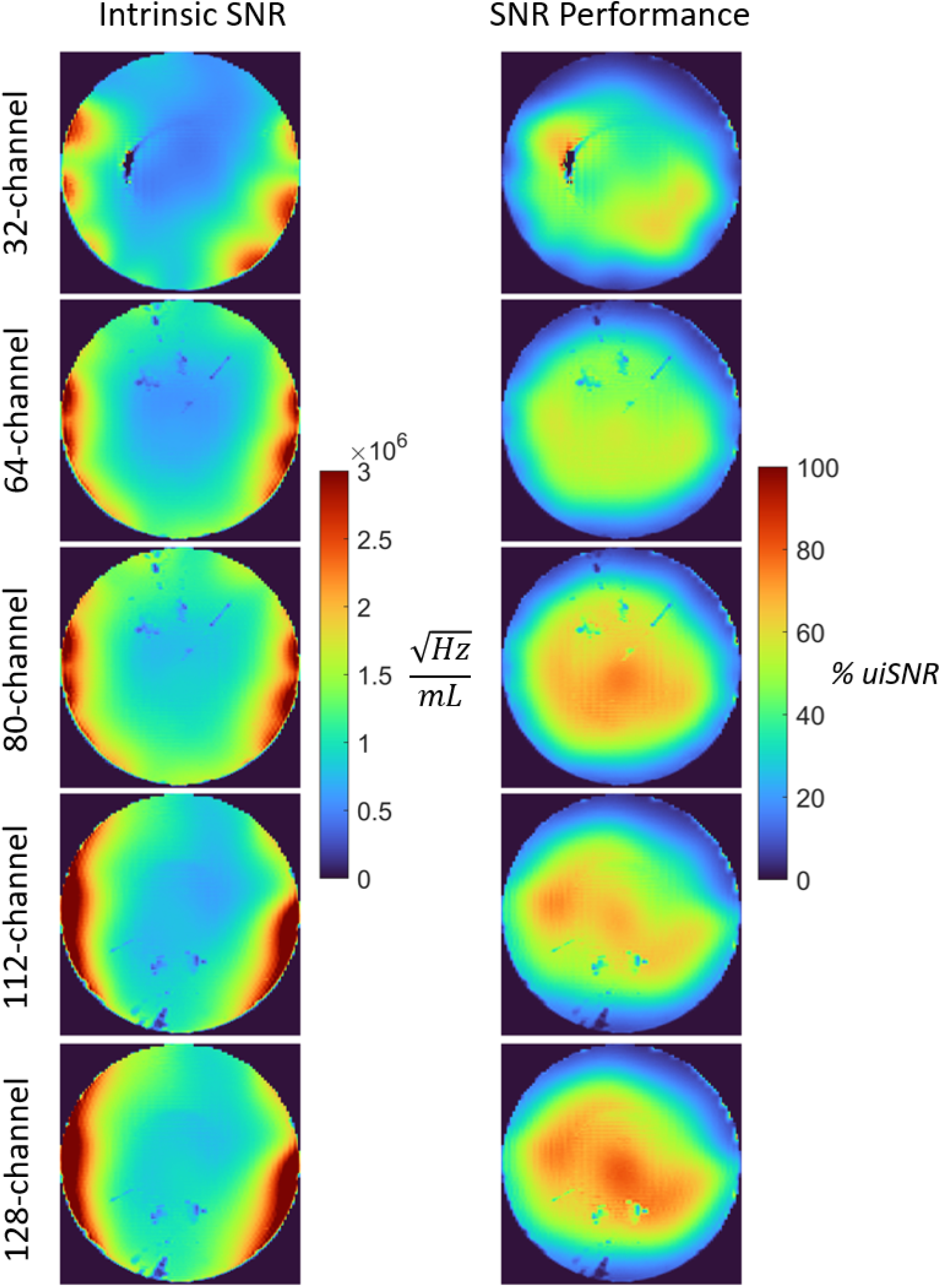
Intrinsic SNR maps (left) and uiSNR-normalized maps (right) for a 32-channel (Nova) and 64-channel (NeuroCam) coil at 7T.

To summarize these observations in a more compact, depth-resolved form, Figure 7 shows the intrinsic SNR performance as a function of depth from the phantom surface for all coils under investigation. By averaging SNR within shell-shaped regions of interest at increasing depths. In addition, to quantify the SNR gain with B_0_ as a function of depth within the phantom, the best-performing coil at each field strength was selected at each depth for curve fitting (see Supporting Figure S1).

**Figure 7:**
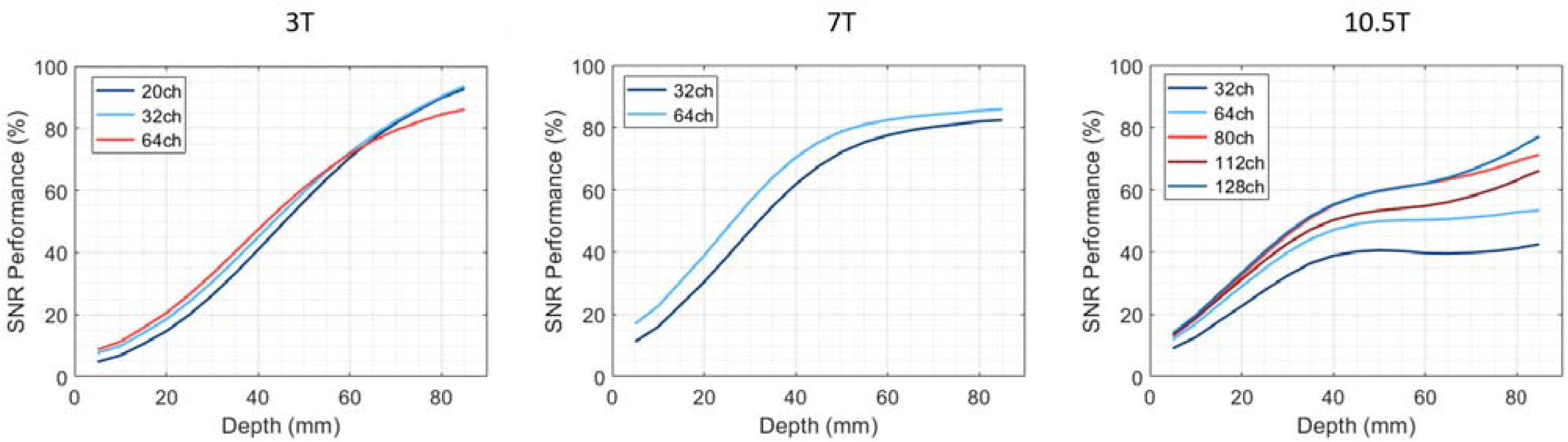
Plots of intrinsic SNR performance as a function of depth from the phantom surface for all coils under investigation. SNR is averaged within shell-shaped regions of interest at increasing depths to generate these plots.

### 3.2 Quantification of Parasitic Losses

During Q-factor measurements, a variety of decoupled probes were compared to determine the bias of the probe. Unloaded Q is underestimated by up to 17% with an improperly decoupled B-field probe (S_21_ = -35 dB) vs the properly decoupled probe described in methods (S_21_ = -65 dB). A properly decoupled B-field probe is essential and all measurements detailed below are made with maximal possible decoupling.

Figure 8 presents measured (solid line) and simulated (dashed line) Q-ratios for 7T and 10.5T loop resonators ranging in diameter between 2 cm and 8 cm. Plots show Q-ratio as a function of sample distance ranging from 1 cm to 5 cm. This same data is plotted again in terms of ratio of SNR over SNR_ideal_ for 7T and 10.5T.

**Figure 8:**
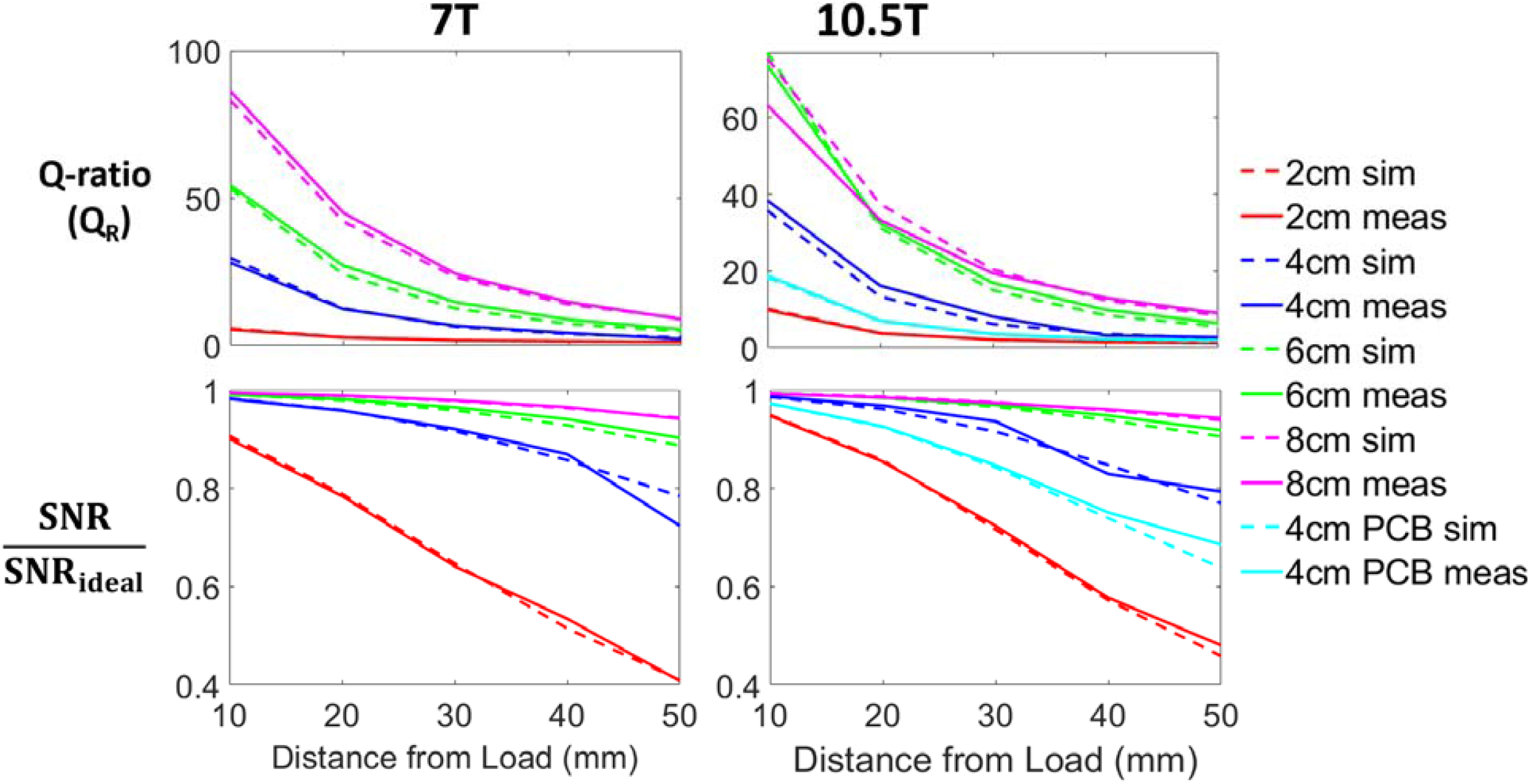
Plots of Q-ratio as a function of sample distance for measured (solid line) and simulated (dashed line) Q-ratios for 7T (left) and 10.5T (right) loop resonators ranging in diameter between 2 cm and 8 cm. Plotted below is the same data in terms of ratio of SNR over SNR_ideal_ for 7T and 10.5T.

Radiation resistance calculated from EM simulations for the unloaded and loaded resonators is plotted in Supporting Figure S2A relative to coil diameter for 7T (blue plot) and 10.5T (red plot). These data are also plotted as a ratio in Supporting Figure S2B. In this plot, the ratio of radiation resistance in the unloaded case vs the loaded case is shown as a function of distance from the load. Supporting Figure S3 separates out coil resistance, sample resistance, and radiation resistance as calculated from simulation as a function of field strength and distance to the load.

### 3.3 Bench-measurement-informed Receive Arrays Simulation

Table 1 summarizes the hybrid technique used to quantify parasitic losses in individual loops of the 128-channel coil by combining bench Q-ratio measurements with single-loop simulations. The table presents the breakdown of parasitic-loss components at different sample-to-load distances. In addition, to highlight the implications of these parasitic losses for sample-noise dominance and, ultimately, SNR, the corresponding noise power ratios (NPRs), defined as *R*_*parasitic*_/R_*sampie*_), are shown in Supporting Figure S4.

**Table 1.** Breakdown of parasitic-loss components (Ω) for representative loop geometries at different sample-to-coil distances.

| Coil geometry | Distance (mm) | Sample | Conductor | Radiation | Capacitors | Matching circuit | Detuning circuit |
| --- | --- | --- | --- | --- | --- | --- | --- |
| 4 cm diameter circular | 10 | 4.687 | 0.280 | 0.050 | 0.532 | 0.010 | 1.343 |
|  | 20 | 1.862 |  | 0.068 |  | 0.058 | 1.321 |
|  | 30 | 0.855 |  | 0.085 |  | 0.107 | 1.054 |
|  | 40 | 0.437 |  | 0.102 |  | 0.158 | 0.541 |
| 5 × 5 cm<sup>2</sup> square | 10 | 13.778 | 0.460 | 0.215 | 0.634 | 0.259 | 2.941 |
|  | 20 | 6.057 |  | 0.283 |  | 1.239 | 2.067 |
|  | 30 | 2.953 |  | 0.349 |  | 1.967 | 1.661 |
|  | 40 | 1.582 |  | 0.411 |  | 2.443 | 1.724 |
| 7 × 4 cm<sup>2</sup> elliptical | 10 | 14.274 | 0.496 | 0.277 | 0.713 | 0.089 | 2.983 |
|  | 20 | 6.509 |  | 0.360 |  | 0.121 | 2.686 |
|  | 30 | 3.266 |  | 0.438 |  | 0.182 | 2.219 |
|  | 40 | 1.790 |  | 0.514 |  | 0.273 | 1.581 |
| 4 cm equilateral triangular | 10 | 2.071 | 0.257 | 0.018 | 0.293 | 0.867 | 0.010 |
|  | 20 | 0.777 |  | 0.024 |  | 0.518 | 0.105 |
|  | 30 | 0.343 |  | 0.030 |  | 0.407 | 0.134 |
|  | 40 | 0.171 |  | 0.035 |  | 0.535 | 0.086 |

Figure 9 summarizes the experimental and simulated SNR, as well as the corresponding SNR performance relative to uiSNR, for the 128-channel coil. In Figure 9A, the absolute SNR maps are compared across three cases: experimental measurement, simulation including only conductor and radiation losses, and simulation including the full set of measured parasitic losses. In Figure 9B, the corresponding SNR/uiSNR maps are shown for the same three cases. When only conductor and radiation losses are considered, the simulated central performance reaches 93% of uiSNR. After incorporation of all measured parasitic losses, including matching and detuning circuit losses, the simulated central performance decreases to 78%, in much closer agreement with the experimentally measured value of 77%, previously reported by Lagore et al.^30^

**Figure 9:**
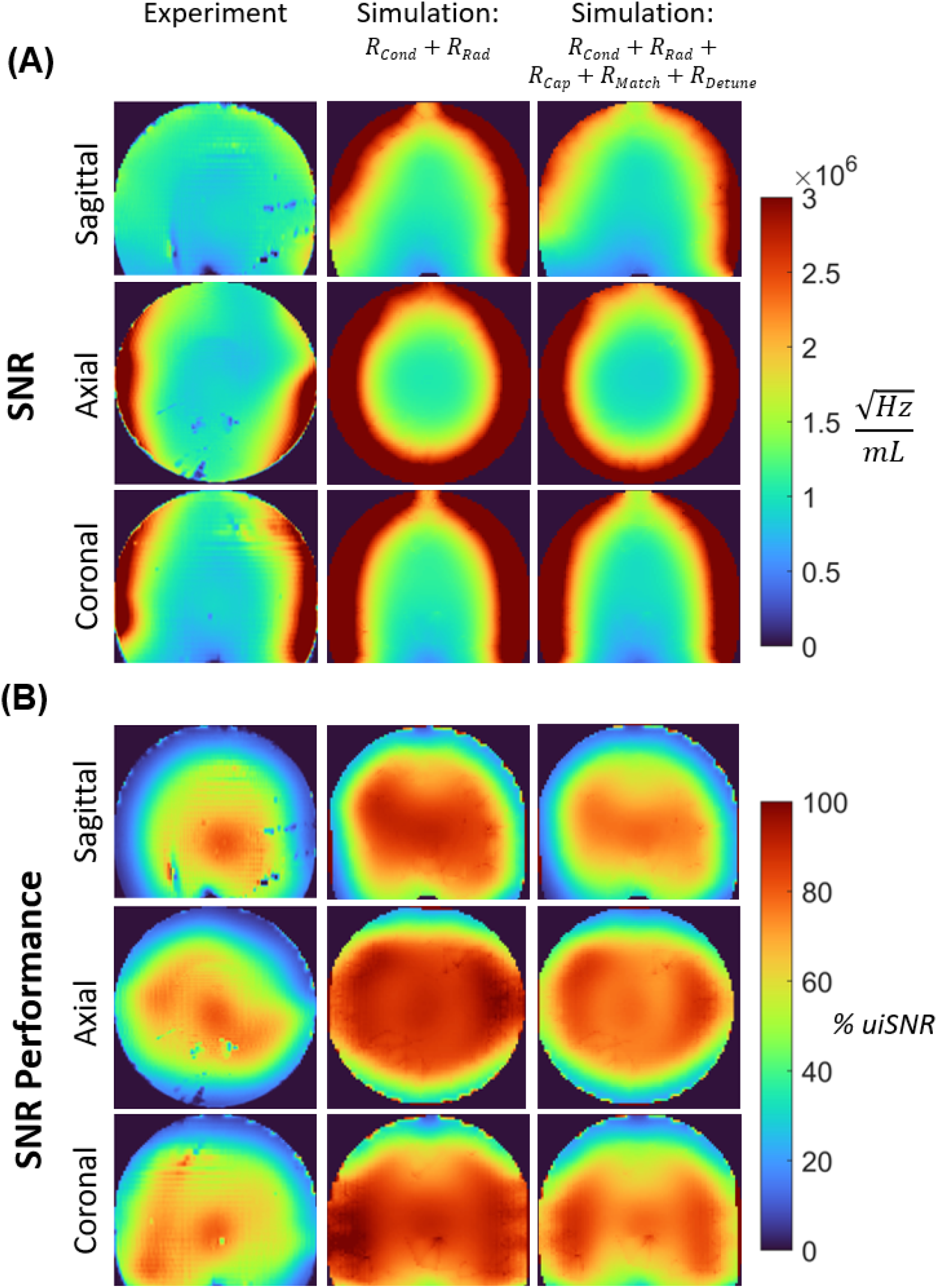
SNR maps of the 128Rx array in the sagittal, axial, and coronal planes (A) acquired experimentally (left column) compared to EM simulations where only R_Cond_ and R_Rad_ areaccounted for (center) vs EM simulations where all losses are accounted for (right). This same data is plotted as a percentage of uiSNR in (B).

## 4 Discussion

### 4.1 SNR Performance Trends Across Field Strength and Coil Architectures

The present results show that the inherently higher SNR available at 10.5T can only be realized when the receive array is designed with much tighter control of practical loss mechanisms than is typically required at lower field strengths. While prior uiSNR-based studies^33^ and coil-specific simulations suggested that high fractions of intermediate and central uiSNR should be achievable with loop arrays, our earlier 10.5T high-channel-count coils fell short of those expectations,^30,31^ particularly in receive-only configurations. The updated results indicate that this shortfall was not due to a failure of the underlying uiSNR framework, but rather to the accumulation of parasitic losses that became increasingly consequential in this UHF regime.

A second important observation (see Supporting Figure S5) is that larger loops at 10.5T— approximately in the size range corresponding to the overlapping 64-channel architecture (∼6cm diameter loops)—appear to be more resilient to parasitic losses than the smaller elements used in our previous 128-channel coil design (∼4cm diameter loops). At the same time, this should not be interpreted as an argument against higher channel count. More channels remain desirable, and necessary, for higher parallel imaging performance. Rather, the present results suggest that loop number, loop size, overlap strategy, and sample-to-coil distance must be optimized together, and that a systematic study will ultimately be needed to identify the best trade-off between intrinsic SNR and accelerated imaging performance at 10.5T.

Another important contribution of this work is the assembly of a rare and informative experimental dataset comprising SNR measurements from multiple commercial and custom-built high–channel-count receive arrays across several field strengths. By acquiring these data within a consistent framework, the variability and bias that often arise when comparing results across the literature and across different centers are largely avoided. This dataset provides direct experimental support for the long-standing prediction that the SNR increases supralinearly with field strength,^45,15^ as shown in Supporting Figure S1. Using the best-performing coil available to us at each field strength at the time of writing this manuscript, we observed that the field-strength dependence of SNR at the center follows an exponent of 1.8, approaching quadratic behavior, while becoming more linear toward the periphery, with an approximate exponent of 1.5 across most intermediate depths.

### 4.2 Q-Ratio and the Quantification of Parasitic Losses at Ultra-High Field

A central methodological point of this study is that unloaded Q must be measured in a way that reflects realistic coil operation at UHF. Conventional unloaded-Q measurements^35,46^ performed in free space include radiation loss, which was historically a small effect in lower-field MRI but becomes substantial at high and ultra-high fields. The problem is not only that radiation lowers unloaded Q, but that the unloaded radiation resistance is not representative of the loaded case. As shown by the simulations, radiation loss is much larger in the unloaded condition than in the loaded condition; thus, including it in the unloaded measurement artificially depresses the Q-ratio and leads to an overestimation of electronic-noise penalty. This is the reason for using the conductive plane during unloaded-Q measurements.^41^ The conductive plane suppresses the unrealistically large, unloaded radiation contribution and yields a more meaningful estimate of the balance between sample noise and the practical electronic losses that remain under realistic loading.

The parasitic-loss breakdown further shows that all of the seemingly “small” loss terms matter. Individually, some of these losses may appear modest, but collectively they can account for a substantial fraction of the missing SNR. This is an important design lesson at 10.5T: performance is not lost through a single catastrophic mechanism, but through the cumulative effects of conductor/substrate loss, capacitor ESR, matching loss, detuning loss, and feed-related implementation details. For this reason, the commonly invoked Q-ratio threshold of 2— regarded as the lower bound of the sample-noise-dominant regime^29,47^ and has served as a key criterion in optimizing loop size^29^—is not sufficient for high-performance receive arrays evaluation at UHF. Even in an idealized case, that threshold still corresponds to a substantial SNR penalty (∼30% loss), and once additional parasitic losses are introduced throughout the full implementation, an even larger fraction of the available SNR is lost. We propose a more appropriate practical target is a final Q-ratio of approximately 5 with the full circuitry in place, corresponding to only about 10% SNR loss, without requiring aggressive increases in loop size or reductions in channel count.

### 4.3 Bench-Informed Simulation Explains the 128-Channel Performance Gap

One of the most important outcomes of this work is that the experimentally observed performance of the 128-channel coil can be reconciled with simulation once the measured parasitic losses are incorporated into the model. In other words, we matched the experimental results in simulation by starting from bench Q-ratio measurements, extracting the corresponding parasitic penalties, and propagating them into the EM-based SNR calculation. This result provides a direct explanation for why the 128-channel coil did not perform as earlier idealized expectations suggested. The issue was not that the physics-based SNR predictions were fundamentally incorrect, but that the conventional simulation model did not include the full burden of practical implementation losses.

In the case of the 128-channel coil, the dominant parasitic contribution was the detuning circuit. For the feed topology used here, the off-state resistance of the PIN diode was identified as the main source of detuning loss, and its effect can be represented analytically as an additional series resistance in the loop. We derived an equation to represent this off-state resistance as a series resistance in the loop which can simply be added to the other resistances in the Q-factor calculation as such:

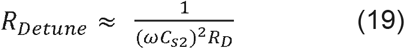

This is a useful result because it points to a clear engineering target: the detuning penalty can be reduced by choosing lower-loss detuning components and/or revisiting the detune-circuit topology itself. Another dominant contributor was the loss associated with the PCB-based loop implementation, which added substrate loss to the underlying conductor loss. This suggests that replacing PCB conductors with lower-loss alternatives, such as high-quality silver-coated copper wire where feasible, could further improve realized SNR. Taken together, these results explain why the 128-channel coil underperformed relative to idealized predictions and show precisely where the design can be improved.

### 4.4 Design Implications for High-Channel-Count Receive Arrays at 10.5T

The main design implication is that, at 10.5T, coil performance is governed by a delicate synergy between loop number, loop size, layout, material choice, component losses, and sample-to-coil distance. This is where the physics meets engineering: uiSNR theory defines what is available in principle, but whether that limit is approached in practice depends on meticulous control of parasitic losses. The present findings therefore argue for a design strategy in which component selection and circuit topology are treated as primary determinants of SNR, rather than as secondary implementation details.

The results also clarify the role of integrated Tx/Rx elements. Supporting Information S6 suggests that when an Rx array is sufficiently low in parasitic loss, it can already meet the numerical uiSNR predictions, and the addition of Tx/Rx elements offers little further SNR benefit. However, when the receive basis lacks sufficiently efficient modes, any additional mode—even if somewhat noise correlated—can help recover performance. This retrospectively justifies the previously proposed strategy of incorporating large Tx/Rx loops into the 80-channel design.^31^ It also suggests that even more efficient gains may be possible by adding truly orthogonal modes, such as dipoles. As demonstrated in Supporting Information S7, preliminary tests in which 7 loops of a 10.5T 32-channel loop array^27^ were replaced with 7 NODES dipoles^48^) yielded highly promising results. Together, these findings support the broader idea that future 10.5T arrays may benefit from hybrid receive bases optimized not only in terms of element number and size, but also for modal complementarity.

### 4.5 Limitations and Future Directions

This study focused on explaining the SNR gap in existing 10.5T receive arrays and on building a practical framework for incorporating parasitic losses into simulation. However, several natural extensions remain. First, a systematic optimization of loop count for accelerated imaging has not yet been performed, and this will likely be required to identify the true performance sweet spot at 10.5T. Second, tighter-fitting, flexible, or conformal loop arrays could reduce sample-to-coil distance and help recover additional SNR, especially if higher channel counts are still desired. Third, future designs should continue to evaluate whether some loop modes are better replaced by more orthogonal elements, such as dipoles, rather than simply adding more loops.

More broadly, the methods presented here offer a practical path for early-stage RF coil development. By combining Q-ratio measurements with validated EM modeling, achievable coil performance can be estimated before a full prototype is finalized. This should be particularly valuable at extreme UHF, where iteration is expensive and where small engineering choices can produce disproportionately large SNR consequences.

## Conclusions

We found that the long-standing gap between predicted and measured SNR performance of high-channel-count 10.5T loop arrays is largely explained by parasitic losses that are not captured in conventional simulations. By quantifying these losses with Q-ratio measurements and incorporating them into EM-based SNR modeling, we were able to reconcile simulation with experiment and identify the dominant engineering penalties limiting coil performance. These results show that, at 10.5T, approaching the uiSNR limit requires not only appropriate loop geometry and channel count, but also systematic minimization of conductor, substrate, component, and feed-circuit losses. This framework provides practical guidance for the design of next-generation high-channel-count receive arrays at UHF.

## References

1. Duyn JH, van Gelderen P, Li TQ et al. High-field MRI of brain cortical substructure based on signal phase. Proc Natl Acad Sci U S A. (2007);104:11796–11801. doi: 10.1073/pnas.0610821104

2. Budde J, Shajan G, Hoffmann J, Uğurbil K, Pohmann R. Human imaging at 9.4 T using T2*-, phase-, and susceptibility-weighted contrast. Magnetic resonance in medicine. (2011);65:544–550.

3. Obusez EC, Lowe M, Oh SH et al. 7T MR of intracranial pathology: Preliminary observations and comparisons to 3T and 1.5T. Neuroimage. (2018);168:459–476. doi:10.1016/j.neuroimage.2016.11.030

4. Sati P. Diagnosis of multiple sclerosis through the lens of ultra-high-field MRI. J Magn Reson. (2018);291:101–109. doi:10.1016/j.jmr.2018.01.022

5. De Martino F, Yacoub E, Kemper V et al. The impact of ultra-high field MRI on cognitive and computational neuroimaging. Neuroimage. (2018);168:366–382. doi:10.1016/j.neuroimage.2017.03.060

6. Dumoulin SO, Fracasso A, van der Zwaag W, Siero JCW, Petridou N. Ultra-high field MRI: Advancing systems neuroscience towards mesoscopic human brain function. Neuroimage. (2018);168:345–357. doi:10.1016/j.neuroimage.2017.01.028

7. Ugurbil K. Imaging at ultrahigh magnetic fields: History, challenges, and solutions. Neuroimage. (2018);168:7–32. doi:10.1016/j.neuroimage.2017.07.007

8. Goense J, Bohraus Y, Logothetis NK. fMRI at High Spatial Resolution: Implications for BOLD-Models. Frontiers in Computational Neuroscience. (2016);10. doi: 10.3389/fncom.2016.00066

9. Huber L, Handwerker DA, Jangraw DC et al. High-Resolution CBV-fMRI Allows Mapping of Laminar Activity and Connectivity of Cortical Input and Output in Human M1. Neuron. (2017);96:1253–1263 e1257. doi:10.1016/j.neuron.2017.11.005

10. Ugurbil K. ULTRAHIGH FIELD and ULTRAHIGH RESOLUTION fMRI. Curr Opin Biomed Eng. (2021);18. doi:10.1016/j.cobme.2021.100288

11. Vaughan JT, Garwood M, Collins CM et al. 7T vs. 4T: RF power, homogeneity, and signal-to-noise comparison in head images. Magnetic Resonance in Medicine: An Official Journal of the International Society for Magnetic Resonance in Medicine. (2001);46:24–30.

12. Wiesinger F, Boesiger P, Pruessmann KP. Electrodynamics and ultimate SNR in parallel MR imaging. Magn Reson Med. (2004);52:376–390. doi:10.1002/mrm.20183

13. Pohmann R, Speck O, Scheffler K. Signal-to-noise ratio and MR tissue parameters in human brain imaging at 3, 7, and 9.4 tesla using current receive coil arrays. Magn Reson Med. (2016);75:801–809. doi:10.1002/mrm.25677

14. Guerin B, Villena JF, Polimeridis AG et al. The ultimate signal-to-noise ratio in realistic body models. Magn Reson Med. (2017);78:1969–1980. doi:10.1002/mrm.26564

15. Le Ster C, Grant A, Van de Moortele PF et al. Magnetic field strength dependent SNR gain at the center of a spherical phantom and up to 11.7T. Magn Reson Med. (2022);88:2131–2138. doi:10.1002/mrm.29391

16. Sadeghi-Tarakameh A, DelaBarre L, Lagore RL et al. In vivo human head MRI at 10.5T: A radiofrequency safety study and preliminary imaging results. Magn Reson Med. (2020);84:484–496. doi:10.1002/mrm.28093

17. Hingerl L, Strasser B, Schmidt S et al. Exploring In Vivo Human Brain Metabolism at 10.5 T: Initial Insights from MR Spectroscopic Imaging; Initial Insights from MR Spectroscopic Imaging. Neuroimage. (2024);Under Revision.

18. Vizioli L, Moeller S, Dowdle L et al. Spanning spatial scales with functional imaging in the human brain; initial experiences at 10.5 Tesla. bioRxiv. (2024);2024–2012.

19. Qu SX, Liu J, van Gelderen P et al. Advancing whole-brain BOLD functional MRI in humans at 10.5 T with motion-robust 3D echo-planar imaging, parallel transmission, and high-density radiofrequency receive coils. Magnetic Resonance in Medicine. (2026);95:1068–1088. doi:10.1002/mrm.70110

20. Ocali O, Atalar E. Ultimate intrinsic signal-to-noise ratio in MRI. Magn Reson Med. (1998);39:462–473. doi:10.1002/mrm.1910390317

21. Lattanzi R, Sodickson DK. Ideal current patterns yielding optimal signal-to-noise ratio and specific absorption rate in magnetic resonance imaging: computational methods and physical insights. Magn Reson Med. (2012);68:286–304. doi:10.1002/mrm.23198

22. Wiggins GC, Triantafyllou C, Potthast A et al. 32-channel 3 Tesla receive-only phased-array head coil with soccer-ball element geometry. Magn Reson Med. (2006);56:216–223. doi:10.1002/mrm.20925

23. Schmitt M, Potthast A, Sosnovik DE et al. A 128-channel receive-only cardiac coil for highly accelerated cardiac MRI at 3 Tesla. Magn Reson Med. (2008);59:1431–1439. doi:10.1002/mrm.21598

24. Wiggins GC, Polimeni JR, Potthast A et al. 96-Channel receive-only head coil for 3 Tesla: design optimization and evaluation. Magn Reson Med. (2009);62:754–762. doi:10.1002/mrm.22028

25. Keil B, Blau JN, Biber S et al. A 64-channel 3T array coil for accelerated brain MRI. Magn Reson Med. (2013);70:248–258. doi:10.1002/mrm.24427

26. Ugurbil K, Auerbach E, Moeller S et al. Brain imaging with improved acceleration and SNR at 7 Tesla obtained with 64-channel receive array. Magn Reson Med. (2019);82:495–509. doi:10.1002/mrm.27695

27. Tavaf N, Lagore RL, Jungst S et al. A self-decoupled 32-channel receive array for human-brain MRI at 10.5 T. Magn Reson Med. (2021);86:1759–1772. doi:10.1002/mrm.28788

28. May MW, Hansen SJD, Mahmutovic M et al. A patient-friendly 16-channel transmit/64-channel receive coil array for combined head-neck MRI at 7 Tesla. Magn Reson Med. (2022);88:1419–1433. doi:10.1002/mrm.29288

29. Gruber B, Stockmann JP, Mareyam A et al. A 128-channel receive array for cortical brain imaging at 7 T. Magn Reson Med. (2023);90:2592–2607. doi:10.1002/mrm.29798

30. Lagore RL, Sadeghi-Tarakameh A, Grant A et al. A 128-channel receive array with enhanced signal-to-noise ratio performance for 10.5T brain imaging. Magn Reson Med. (2025);93:2680–2698. doi:10.1002/mrm.30476

31. Waks M, Lagore RL, Auerbach E et al. RF coil design strategies for improving SNR at the ultrahigh magnetic field of 10.5T. Magn Reson Med. (2025);93:873–888. doi:10.1002/mrm.30315

32. Lattanzi R, Grant AK, Polimeni JR et al. Performance evaluation of a 32-element head array with respect to the ultimate intrinsic SNR. NMR Biomed. (2010);23:142–151. doi:10.1002/nbm.1435

33. Vaidya MV, Sodickson DK, Lattanzi R. Approaching Ultimate Intrinsic SNR in a Uniform Spherical Sample with Finite Arrays of Loop Coils. Concepts Magn Reson Part B Magn Reson Eng. (2014);44:53–65. doi:10.1002/cmr.b.21268

34. Zhang B, Radder J, Giannakopoulos I et al. Performance of receive head arrays versus ultimate intrinsic SNR at 7 T and 10.5 T. Magn Reson Med. (2024);92:1219–1231. doi:10.1002/mrm.30108

35. Hayes CE, Axel L. Noise performance of surface coils for magnetic resonance imaging at 1.5 T. Medical physics. (1985);12:604–607.

36. Kumar A, Edelstein WA, Bottomley PA. Noise figure limits for circular loop MR coils. Magn Reson Med. (2009);61:1201–1209. doi:10.1002/mrm.21948

37. Roemer PB, Edelstein WA, Hayes CE, Souza SP, Mueller OM. The NMR phased array. Magn Reson Med. (1990);16:192–225. doi:10.1002/mrm.1910160203

38. Pruessmann KP, Weiger M, Scheidegger MB, Boesiger P. SENSE: sensitivity encoding for fast MRI. Magn Reson Med. (1999);42:952–962. doi:10.1002/(SICI)1522-2594(199911)42:5<952::AID-MRM16>3.0.CO;2-S

39. Keil B, Wald LL. Massively parallel MRI detector arrays. J Magn Reson. (2013);229:75–89. doi:10.1016/j.jmr.2013.02.001

40. Darrasse L, Kassab G. Quick measurement of NMR-coil sensitivity with a dual-loop probe. Review of Scientific Instruments. (1993);64:1841–1844. doi:10.1063/1.1144020

41. Chen G, Collins CM, Sodickson DK, Wiggins GC. A method to assess the loss of a dipole antenna for ultra-high-field MRI. Magn Reson Med. (2017). doi:10.1002/mrm.26777

42. Edelstein WA, Glover GH, Hardy CJ, Redington RW. The intrinsic signal-to-noise ratio in NMR imaging. Magn Reson Med. (1986);3:604–618.

43. Bosma H. On the theory of linear noisy systems, Technische Hogeschool Eindhoven, (1967).

44. Gapais P-F, Luong M, Amadon A. Revisiting the Impact of Inter-Channel Coupling and Thermal Noise Correlation on MRI Receive-Array Performance: A Simulation Study. IEEE Journal of Electromagnetics, RF and Microwaves in Medicine and Biology. (2024);PP:1–8. doi:10.1109/JERM.2024.3509589

45. Pfrommer A, Henning A. The ultimate intrinsic signal-to-noise ratio of loop- and dipole-like current patterns in a realistic human head model. Magn Reson Med. (2018);80:2122–2138. doi:10.1002/mrm.27169

46. Kumar A, Edelstein WA, Bottomley PA. Noise figure limits for circular loop MR coils. Magnetic Resonance in Medicine. (2009);61:1201–1209. doi:10.1002/mrm.21948

47. Webb A, O’Reilly T. Tackling SNR at low-field: a review of hardware approaches for point-of-care systems. Magma. (2023);36:375–393. doi:10.1007/s10334-023-01100-3

48. Sadeghi-Tarakameh A, Waks M, Grant A et al. Boosting Central Head SNR at 10.5T: 32-channel Hybrid RF Coil Comprised of 25 Rx-only Loops and 7 TxRx NODES Dipoles. Proc Int Soc Mag Reson Med. (2023);3913.

