## Supplementary Information for "SNR Enhancement Considerations for Loop Receive Coils at Ultra-High Fields"

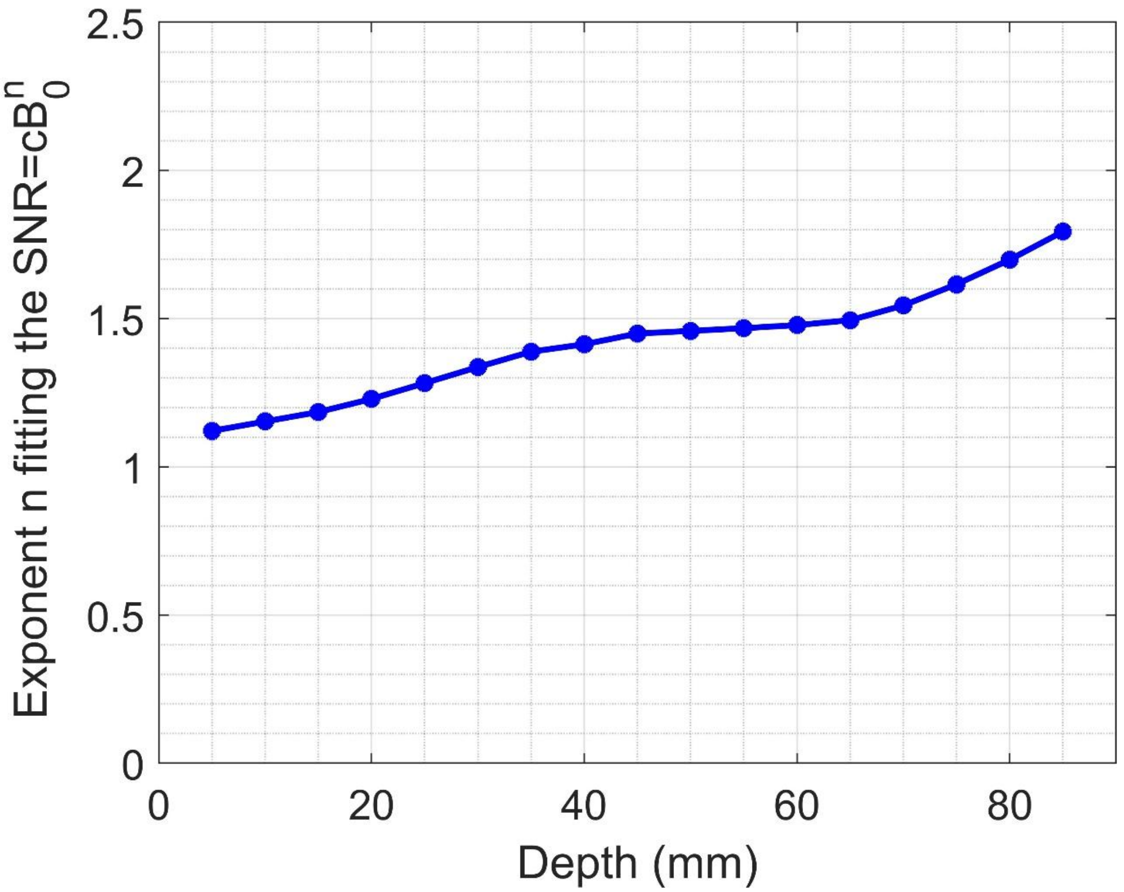


**Figure S1:** Intrinsic SNR performance for the best-performing coil at each field strength to quantify the SNR gain with B0 as a function of depth.

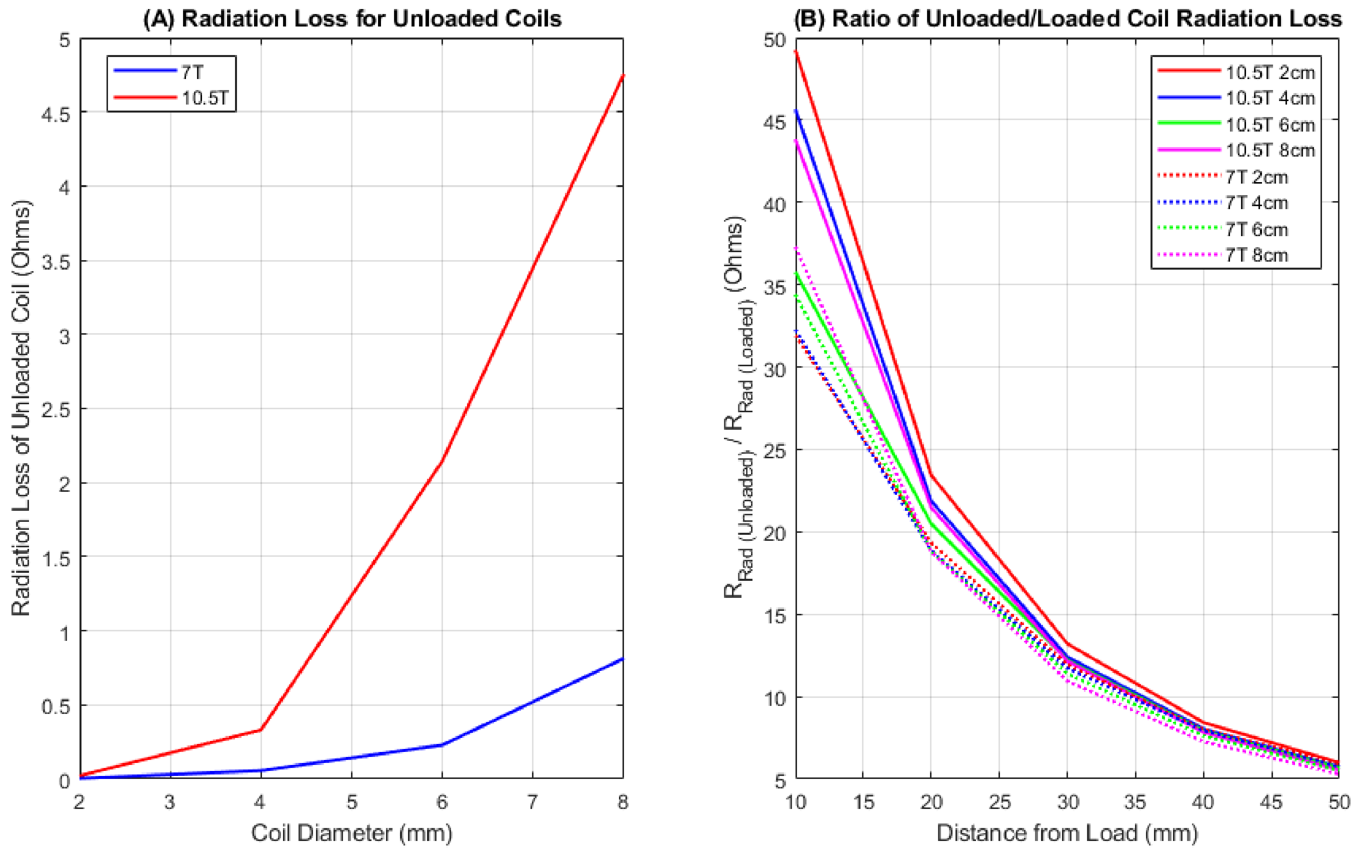


**Figure S2:** Radiation resistance calculated from EM simulations for the unloaded and loaded resonators (A) relative to coil diameter for 7T (blue plot) and 10.5T (red plot). These data are also plotted as a ratio (B). The ratio of radiation resistance in the unloaded case vs the loaded case is shown as a function of distance from the load.

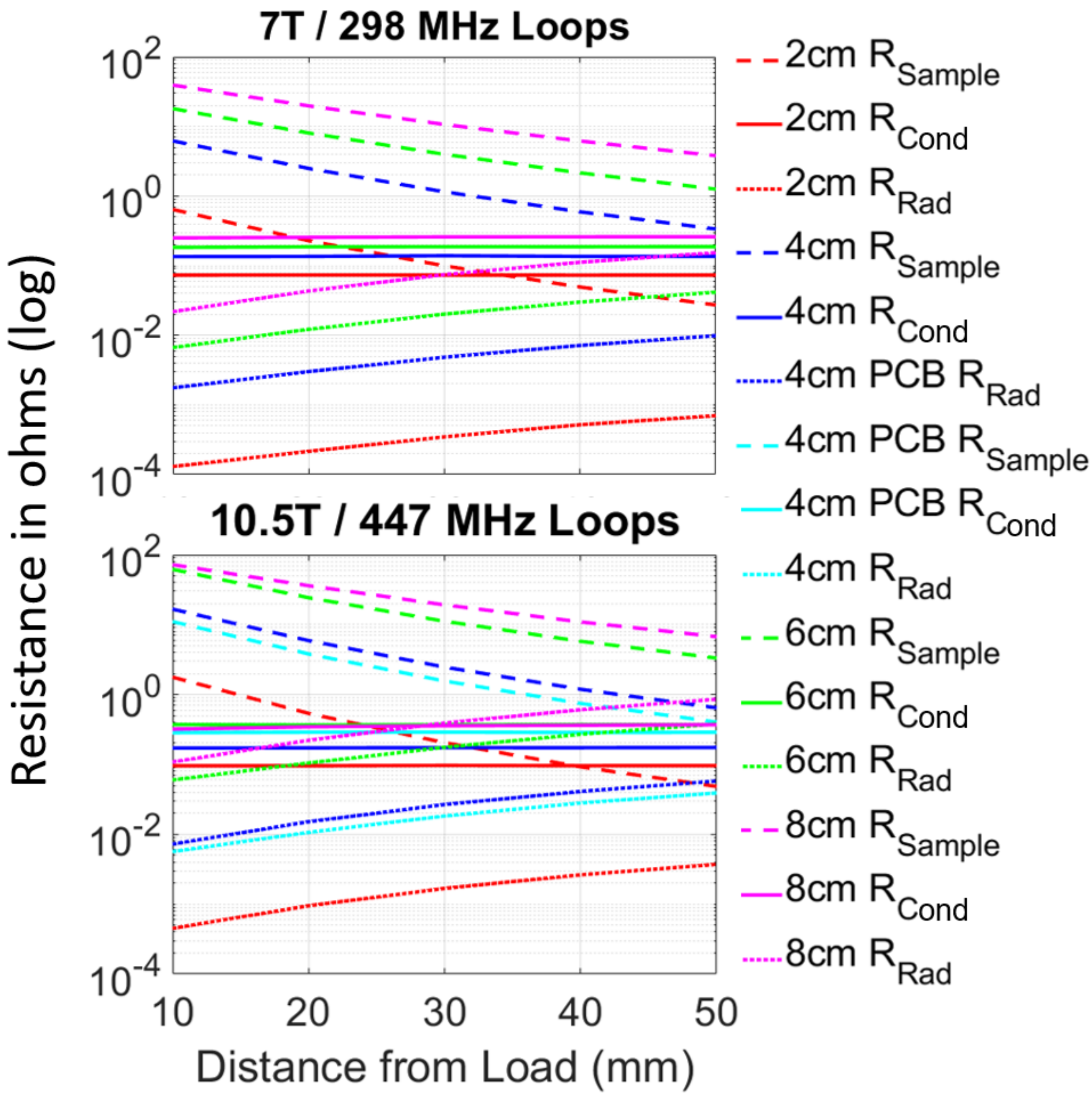


**Figure S3:** Coil resistance ($R_{coil}$), sample resistance ($R_{Sample}$), and radiation resistance ($R_{Rad}$) as calculated from simulation as a function of field strength and distance to the load.

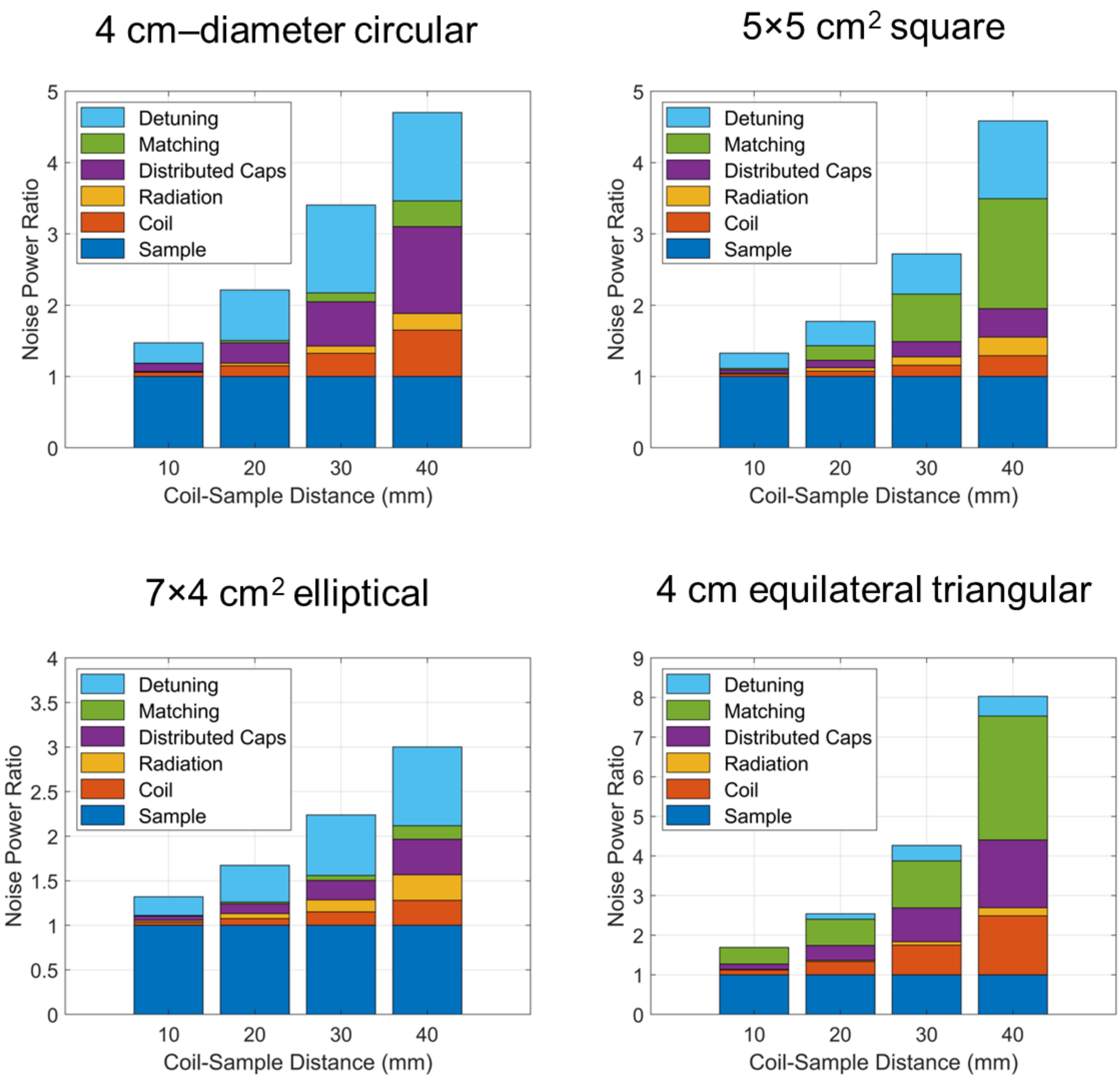


**Figure S4:** A breakdown of parasitic-loss components at different sample-to-load distances in terms of noise power ratios (NPRs), defined as $R_{parasitic}/R_{sample}$ , for different types of loops in the 128Rx array. Note the vertical axis is scaled differently for each of the bottom plots.

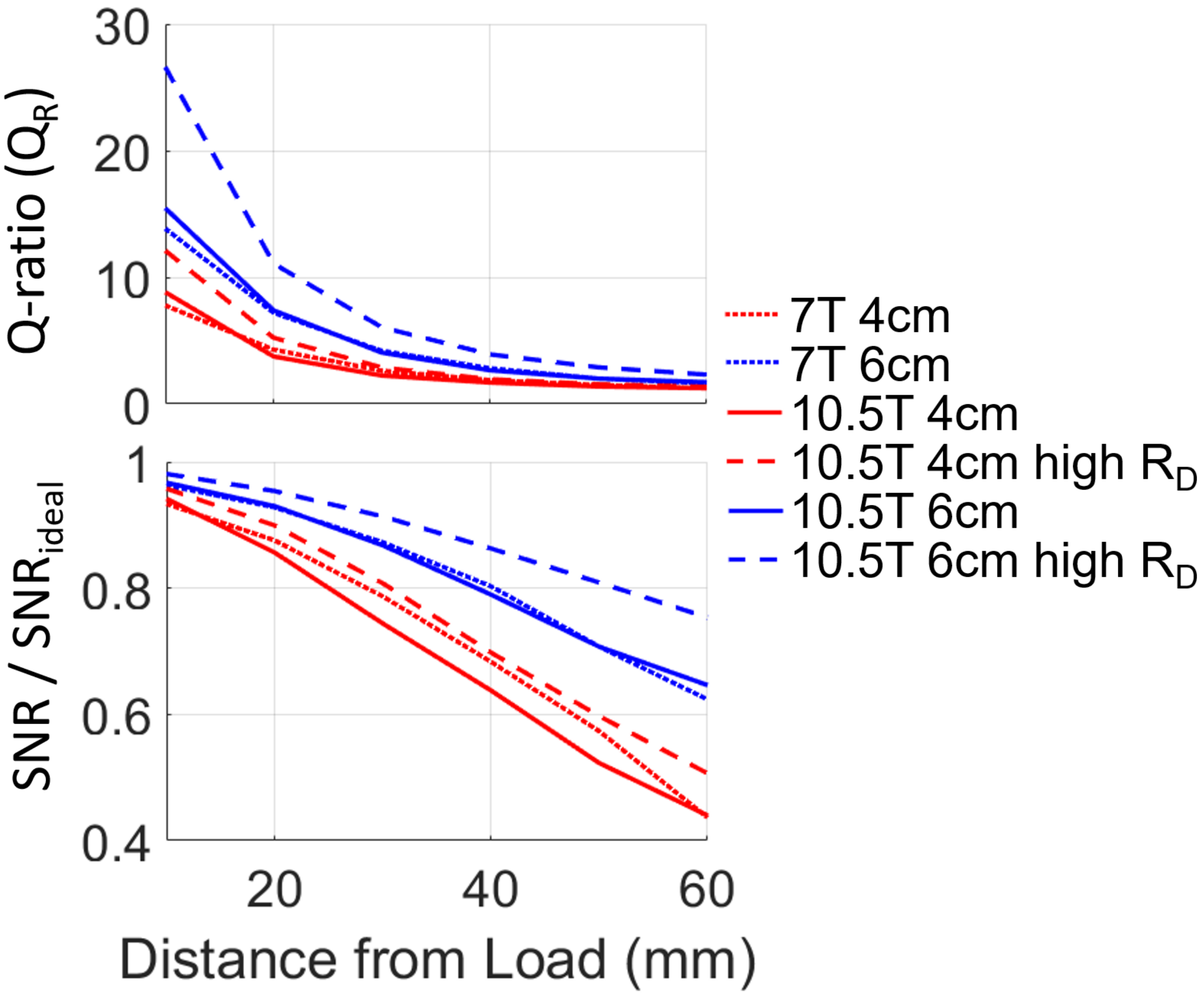


**Figure S5:** (A) Plots practically achievable Q-ratios for RF coils of 4 cm (red) and 6 cm (blue) diameter at 7T (dotted) and 10.5T (dashed and solid) as a function of loop distance from load. In (B) the same data is plotted as a fraction of the SNR captured from a lossless coil. These coils utilize the feed circuit described in Figure 2C with equivalent circuit shown in Figure 2D.

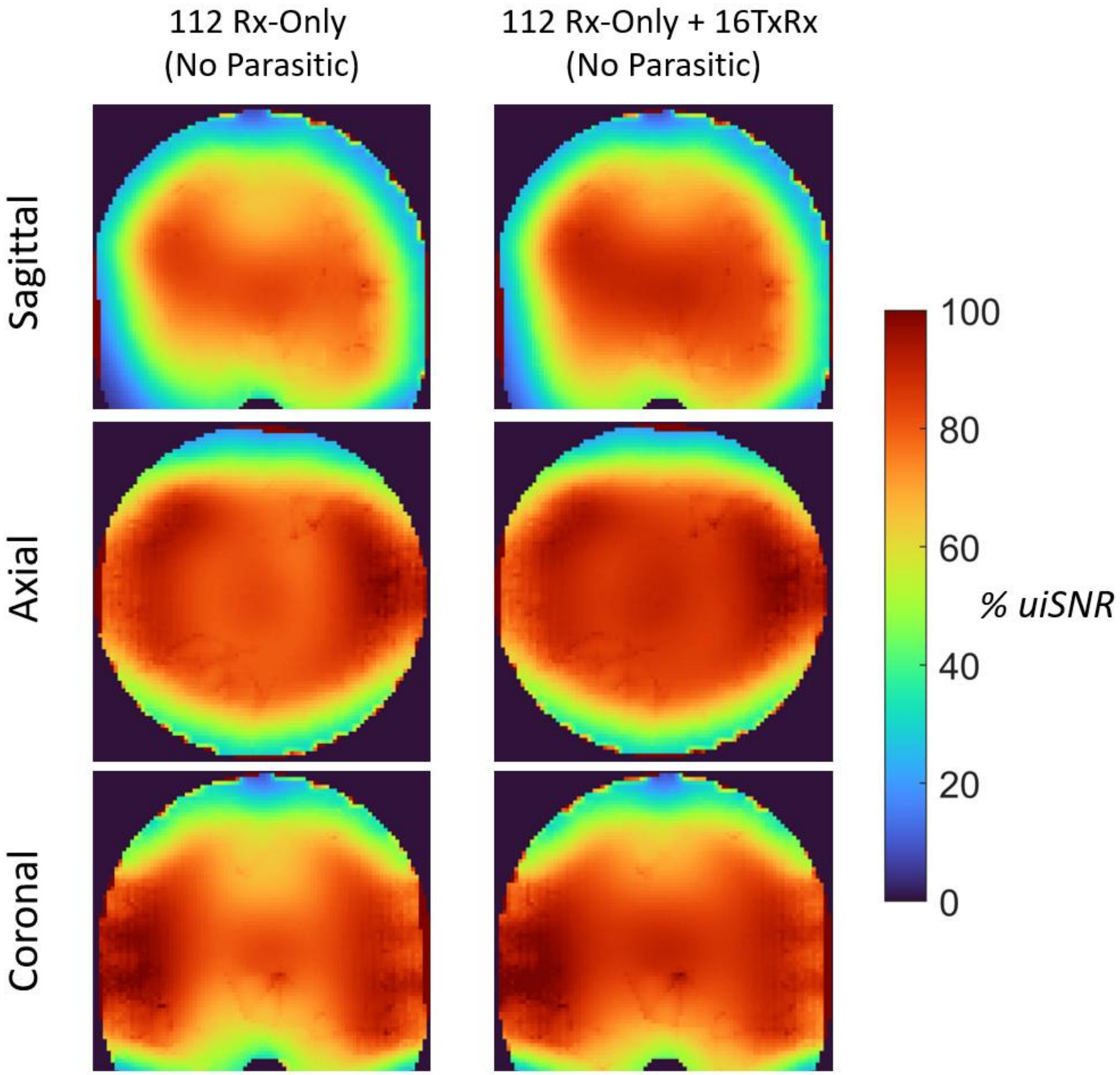


**Figure S6:** SNR as a percentage of uiSNR maps in the sagittal, axial, and coronal planes for the 112 channels of the Rx-only loop array vs the full 128Rx including the 16TxRx elements. There is little additional benefit from the 16TxRx elements when there are no parasitic losses in the 112Rx array.

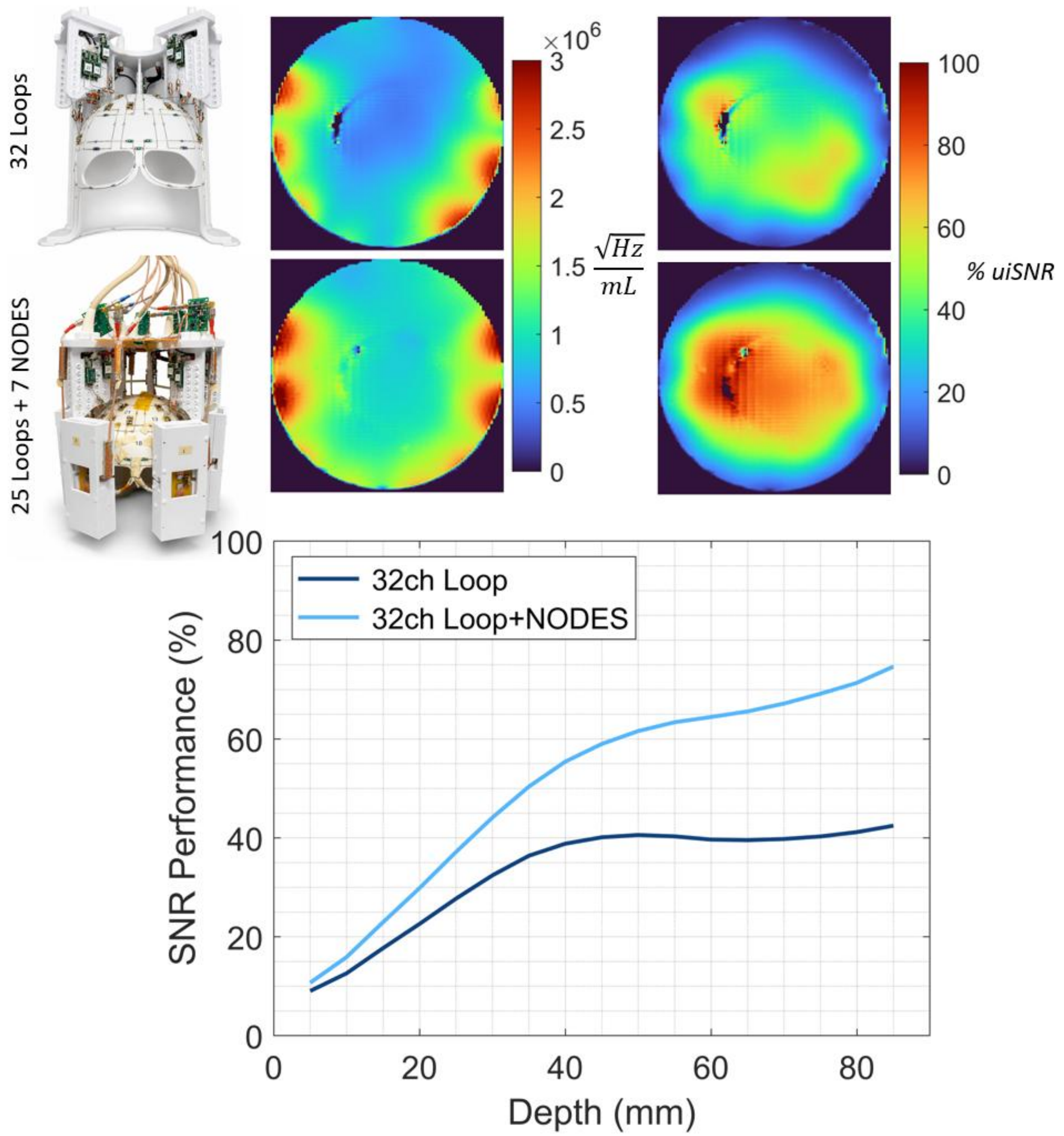


**Figure S7:** Comparison of the 32-channel loop receive array to the 25 loop + 7NODES dipole receive array in terms of SNR maps in the axial slice and percentage of uiSNR with the same data plotted as shells of increasing depth inside the lightbulb-shaped phantom below.
